# A CLONAL LEGACY? REPRODUCTIVE MODE VARIATION IN HARD AND SOFT BOTTOM *GRACILARIA VERMICULOPHYLLA* POPULATIONS

**DOI:** 10.1101/2025.06.19.660572

**Authors:** Alexis P. Oetterer, Solenn Stoeckel, Will H. Ryan, Stacy A. Krueger-Hadfield

## Abstract

The red macroalga *Gracilaria vermiculophylla* invasion provides an opportunity to investigate the evolution of biphasic life cycles and of reproductive modes by understanding how they structure and contribute colonizing new environments in natural conditions. In hard bottom habitats, we find gametophytes and tetrasporophytes fixed by holdfasts to hard substrates, whereas in soft bottom habitats, we find free-living tetrasporophytes either drifting or anchored by tube-building polychaetes. We collected thalli from hard and soft bottom habitats along the Eastern Shore of Virginia and Maryland to investigate the role of substrate on life cycle and reproductive mode dynamics. We determined the phase and sex using observable reproductive structures and a sex-linked PCR assay, followed by genotyping all thalli using nine microsatellite loci. Sexual reproduction prevailed in hard bottom sites, whereas clonal (asexual) reproduction dominated soft bottom sites and was accompanied by tetrasporophytic dominance. There was site-specific variation in selfing and clonal rates that are supported by observations of physiological stress and local extirpation, such as at Ape Hole Creek and Fowling Point, respectively. We found evidence of isolation by distance and the structuring of genetic diversity by habitat type, then site, and finally by year. While broad patterns have been described across the extant range, we clarify population genetic patterns in hard versus soft bottom habitats that are not confounded by the invasion history comparing native and non-native thalli. These results have implications for the on-going spread of this alga and contribute to our understanding of the population genetics of partially clonal taxa.

## INTRODUCTION

The relative rates of sexual and clonal reproduction (i.e., the reproductive mode, Barrett, 2010) are integral to understanding the partitioning of genetic diversity within and among populations (Hamrick & Godt, 1996). Though mixed mating – variable rates of self-fertilization (i.e., selfing) and outcrossing – may be overestimated (Meyer et al., 2024), our knowledge of reproductive mode dynamics has largely focused on hermaphroditic (or monoecious) angiosperms and the rates of selfing and outcrossing (e.g., Barrett et al., 2014; Whitehead et al., 2018). By comparison, we know much less about the rates of clonal reproduction in natural populations, and this is further compounded by a lack of standardization in calculating and reporting population genetics summary statistics (see Arnaud-Haond et al., 2007; Stoeckel et al., 2021a). Partially clonality is common across eukaryotes (Schön et al., 2009) and the relative rates of sexual versus clonal reproduction in a population can affect evolutionary responses to fluctuating environmental conditions (Orive et al., 2017). Empirical data, encapsulating eukaryotic diversity beyond flowering plants, are critical to ultimately understand responses to climate change.

Eukaryotes display tremendous life cycle variation (see Bell, 1994; Beukeboom & Perrin, 2014), rendering a broader synthesis of reproductive mode variation challenging (Krueger-Hadfield, 2024). Predictions based on diploid life cycles may not be readily transferred to other life cycle types in which the haploid phase is dominant or is of long duration (e.g., Krueger Hadfield & Hoban, 2016; Stoeckel et al., 2021a). For example, in diploid life cycles^1^, selfing is only possible in hermaphroditic or monoecious taxa. Yet, in haploid-diploid life cycles with separate sexes (termed dioicous because sex is determined in the haploid phase; Beukeboom & Perrin, 2014), intergametophytic selfing is possible when a male and female gametophytic pair share the same sporophytic parent (Klekowski, 1969). Separate sexes are often used as a proxy to assume obligate outcrossing in angiosperms (see discussion in Krueger-Hadfield et al., 2024) but cannot be used as such in haploid-diploid taxa in which selfing occurs despite cross-fertilization of gametes produced by male and female gametophytes.

Intergametophytic selfing has been observed in mosses (e.g., Eppley et al., 2007) and macroalgae (e.g., Heiser et al., 2023; Krueger-Hadfield et al., 2013, 2015). Moreover, for haploid-diploid taxa, clonal reproduction will result in the loss of a phase in the life cycle. ‘Recycling’ of one phase through clonal reproduction has resulted in gametophyte-only populations in ferns (e.g., Pinson et al., 2017) and sporophytic dominance in macroalgae (e.g., Gabrielsen et al., 2002; Guillemin et al., 2008; Hwang et al., 2005; Krueger-Hadfield et al., 2016; Sosa et al., 1998; Steen et al., 2019; Williams et al., 2024). The recovery of cycling between gametophytes and sporophytes largely depends on which phase is maintained by clonal reproduction (see Krueger-Hadfield, 2020). Haploid-diploid taxa present unique challenges in the exploration of their genetic structure using population genetic tools due to their life cycles (de Meeûs et al., 2007; Krueger-Hadfield, 2020, 2024; Valero et al., 2001) but nevertheless require these tools to determine the prevailing mode of reproduction (Tibayrenc & Ayala, 1991).

There are only a handful of studies that have used genetic markers, such as microsatellites, to investigate reproductive mode dynamics in haploid-diploid macroalgae (reviewed in Krueger-Hadfield, 2024; Krueger-Hadfield et al., 2021a). Across the three evolutionarily diverse macroalgal lineages, there is tremendous reproductive mode variability (Heesch et al., 2021; Krueger-Hadfield et al., 2024; Olsen et al., 2020; Otto & Marks, 1996). Phycologists have long recognized this diversity from laboratory-based cultures (e.g., Maggs, 1988) and macroalgae have been proposed as model eukaryotes that will help fill in knowledge gaps in our understanding of the evolutionary maintenance of sex (Otto & Marks, 1996; Valero et al., 1992). Recent theoretical developments both for partially clonal taxa (Becheler et al., 2017; Stoeckel et al., 2021b) and haploid-diploid macroalgae (Krueger-Hadfield et al., 2019, 2021; Krueger Hadfield & Hoban, 2016; Stoeckel et al., 2021a) allow us to infer reproductive modes from population genetic diversity and its change over time.

The red algal genus *Gracilaria* Greville has received significant population genetic (e.g., Ayres-Ostrock et al., 2019; Engel et al., 1999, 2004; Guillemin et al., 2008; Krueger-Hadfield et al., 2016) and genomic attention (e.g., Flanagan et al., 2021; Lipinska et al., 2023, 2024).

*Gracilaria gracilis* was one of the first macroalgae to be studied from a population genetic perspective. Contrary to predictions (Otto & Goldstein, 1992; Otto & Marks, 1996), the populations studied were not only predominantly outcrossed (i.e., allogamous) but there were also few repeated tetrasporophytes as would be expected from mitotic copies of the zygote in the carposporophytes (Engel et al., 1999, 2004; Lavaut et al., 2023). *Gracilaria* spp. are also able to clonally propagate through thallus fragmentation, a characteristic that has resulted in the global dominance of this genus for agar production (Kain & Destombe, 1995). Thalli can grow indefinitely and propagate without holdfasts, but to the best of our knowledge thalli do not reattach via secondary attachments or discs (Kain & Destombe, 1995). Krueger-Hadfield et al. (2023) attempted to differentiate between fixed (i.e., attached via a holdfast) and free-living thalli (i.e., drifting with no holdfast) that live in hard versus soft bottom habitats as a starting point to explore unique population dynamics and reproductive modes by site or thallus type in *G. vermiculophylla* (see also Krueger-Hadfield et al., 2025a for further discussion in general for macroalgae). In Chile, Guillemin et al. (2008) documented a shift to clonal reproduction leading to tetrasporophytic dominance because of farmer activity in which thalli were fragmented and ‘planted’ in the mud. This is a very different habitat type from the natural, rocky shore populations of *G. chilensis* in which male and female (haploid) gametophytes alternate with (diploid) tetrasporophytes. The occurrence of free-living thalli is not uncommon, either for this genus or for other red, green, and brown macroalgae (see Norton & Mathieson, 1983), but we still know little of their evolutionary ecology. There are multiple examples of *Gracilaria* spp. inhabiting hard and soft bottom habitats as fixed and free-living thalli, respectively, making the species of this genus excellent models with which to explore reproductive mode dynamics under different ecological conditions.

*Gracilaria vermiculophylla* (Ohmi) Papenfuss (also referred to as *Agarophyton vermiculophyllum* (Ohmi) Gurgel, J.N. Norris, Fredericq) is a prolific invader, affording the opportunity to explore reproductive mode variation during the invasion of novel ecological conditions. Per Baker’s Law, we expected a shift away from sex in the prevailing reproductive mode associated with the colonization of novel habitats where conspecifics may be in limited supply (Baker, 1955; Pannell, 2015; Pannell et al., 2015). *Gracilaria vermiculophylla* is native to the northwestern Pacific but has spread to every temperate coastline in North America, Europe, and northwestern Africa, likely by hitchhiking as packing material with *Magallana gigas* (synonym *Crassostrea gigas*) exports from Japan during the 20^th^ century (Kim et al., 2010; Krueger-Hadfield et al., 2017). The invasion was accompanied by shifts in phenotype (e.g., (Bonthond et al., 2020; Hammann et al., 2016; Sotka et al., 2018) and the prevailing reproductive mode (Krueger-Hadfield et al., 2016), thereby facilitating colonization success. Krueger-Hadfield et al. (2016) documented genetic signatures of sexual reproduction at sites in the native range (Japan, South Korea, and China), where gametophytes and tetrasporophytes were both found. In contrast, in the non-native range (European and North American coasts), signatures of clonal reproduction were common and tetrasporophytes overwhelmingly dominated most sites in which thalli were sampled (Krueger-Hadfield et al., 2016). Many non-native sites were composed of soft sediment without hard substrata in which thalli were free-living whereas native sites had abundant hard substrata with fixed thalli (Krueger-Hadfield et al., 2016). There were some sites in the non-native range where *G. vermiculophylla* thalli were fixed to hard substrata in which cystocarpic female gametophytes and reproductive male gametophytes and tetrasporophytes were observed (Krueger-Hadfield et al., 2016, 2017, 2018). The documented increase in clonal reproduction in the non-native range was likely influenced by the preponderance of invaded soft bottom mudflats in marshes and estuaries. When hard substrate is available, patterns of reproductive mode variation are likely more nuanced than initially described in Krueger-Hadfield et al. (2016).

The native versus non-native dichotomy in sexual versus clonal reproduction is better represented as a contrast between hard and soft bottom habitats which in turn leads to different prevailing reproductive modes (Figure 1). In hard bottom habitats, there is sufficient substratum to provide the necessary ecological conditions for tetrasporic or carposporic settlement and the life cycle can be completed. By contrast, in soft bottom habitats, there is very little or no hard substrata and therefore spores either cannot settle or are immediately abraded on the ebbing or flowing tide. In *Gracilaria vermiculophylla*, this results in consistent tetrasporophytic bias or dominance in both the native and non-native soft bottom habitats (see Krueger-Hadfield et al., 2016, 2017, 2023, 2025b). The dominance of tetrasporophytes is likely due to ecologically relevant phenotypic differentiation between gametophytes and tetrasporophytes (e.g., Krueger Hadfield & Ryan, 2020; Lees et al., 2018; see also in other *Gracilaria* spp. (Destombe et al., 1989, 1992, 1993; Guillemin et al., 2013), though the relevant phenotypes remain enigmatic. Nevertheless, our current understanding of reproductive mode variation in *G. vermiculophylla* is confounded by the invasion history with the preponderance of collections in native, hard bottom habitats versus non-native, soft bottom habitats (Krueger-Hadfield et al., 2016, 2017). The Delmarva Peninsula in the Mid-Atlantic region of the East Coast of the United States (Figure 2) is an exceptional place to explore the consequences of invasion into soft sediment habitats with concomitant consequences on the reproductive mode and haploid-diploid life cycle. Previous work has shown that the sites sampled in this region belong to the same genetic cluster (Krueger-Hadfield et al., 2017) and we can therefore remove the confounding effects of comparing native and non-native populations dynamics while simultaneously exploring site-specific variation based on substratum type (see also Krueger-Hadfield et al., 2025b).

**Figure 1.**
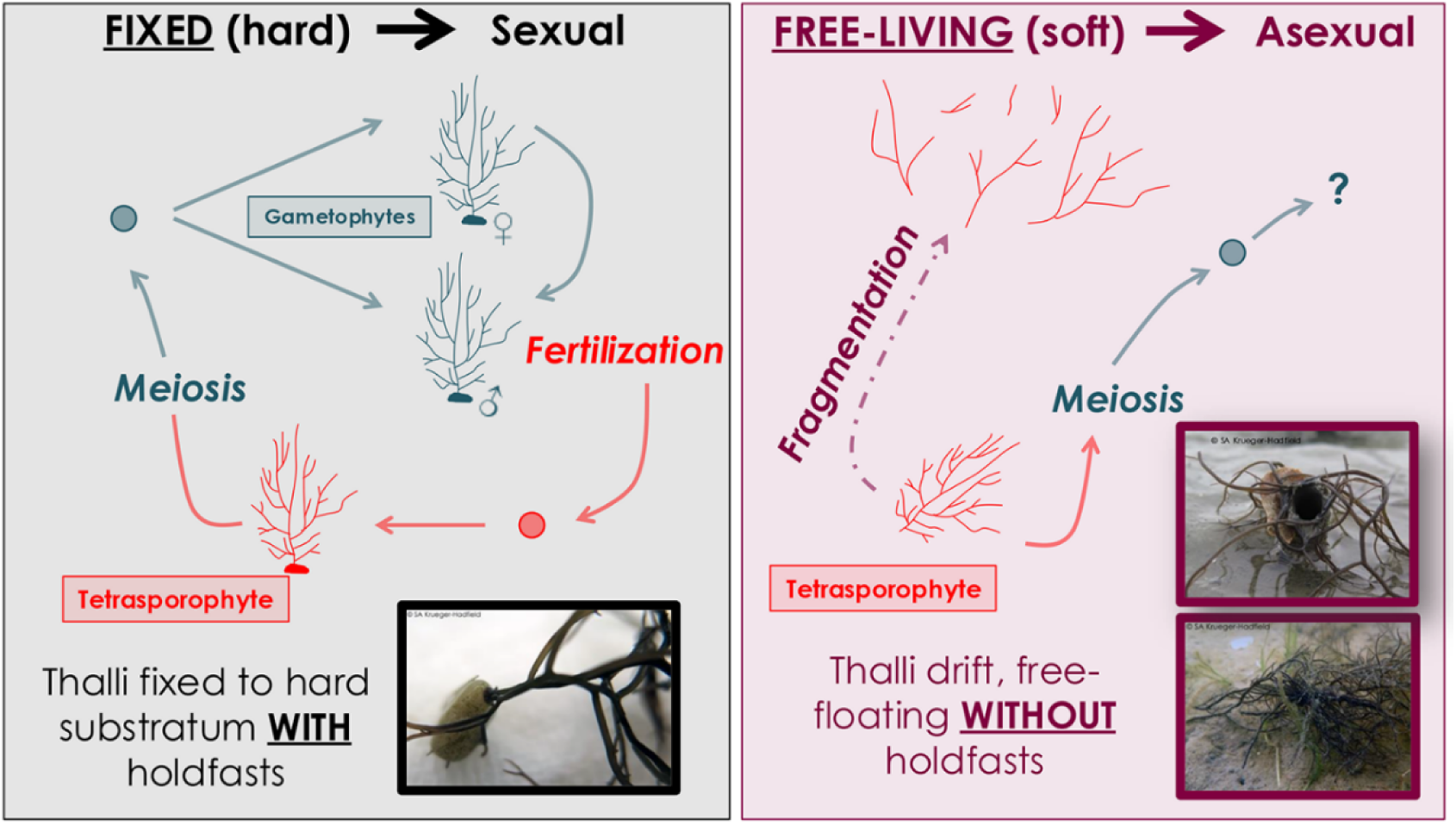
Diagram of the life cycle of fixed versus free-living thalli (thallus size not drawn to scale). In hard bottom habitats, thalli are fixed to hard substrate by a holdfast. Each fixed thallus originates from a spore that settled, germinated, and grew into an adult tetrasporophyte or gametophyte. Hard bottom habitats should be characterized by higher rates of sexual reproduction (see Krueger- Hadfield et al., 2016). We note that the red algal life cycle is simplified here and does not show the third phase – the carposporophytes. In soft bottom habitats, thalli that were once fixed to hard substrate via a holdfast become detached and are free- living or anchored by the decorator polychaete *Dioptra cuprea*. Soft bottom habitats should be characterized by higher rates of clonal reproduction. Animations and photo credits: S.A. Krueger-Hadfield

**Figure 2.**
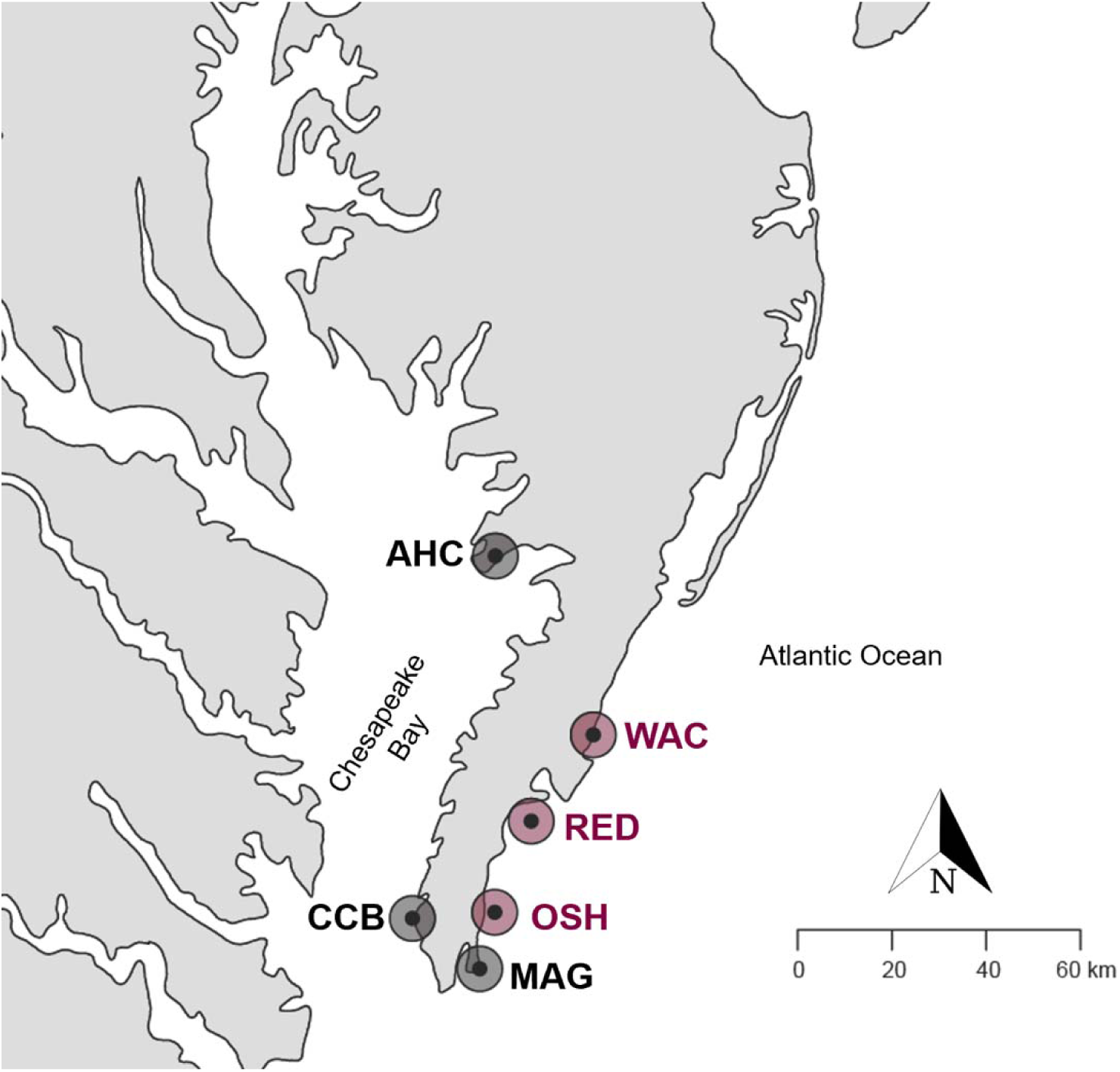
Map of sites sampled for *Gracilaria vermiculophylla* thalli around the Delmarva Peninsula. Colors shown are hard bottom sites with fixed thalli (black) and soft bottom sites with free-living thalli (maroon). AHC, Ape Hole Creek; CCB, Cape Charles Beach; MAG, Magotha Road; OSH, Hillcrest; RED, Fowling Point; WAC, Upper Haul Over. See Table 1 for more detailed site information.

**Table 1.**
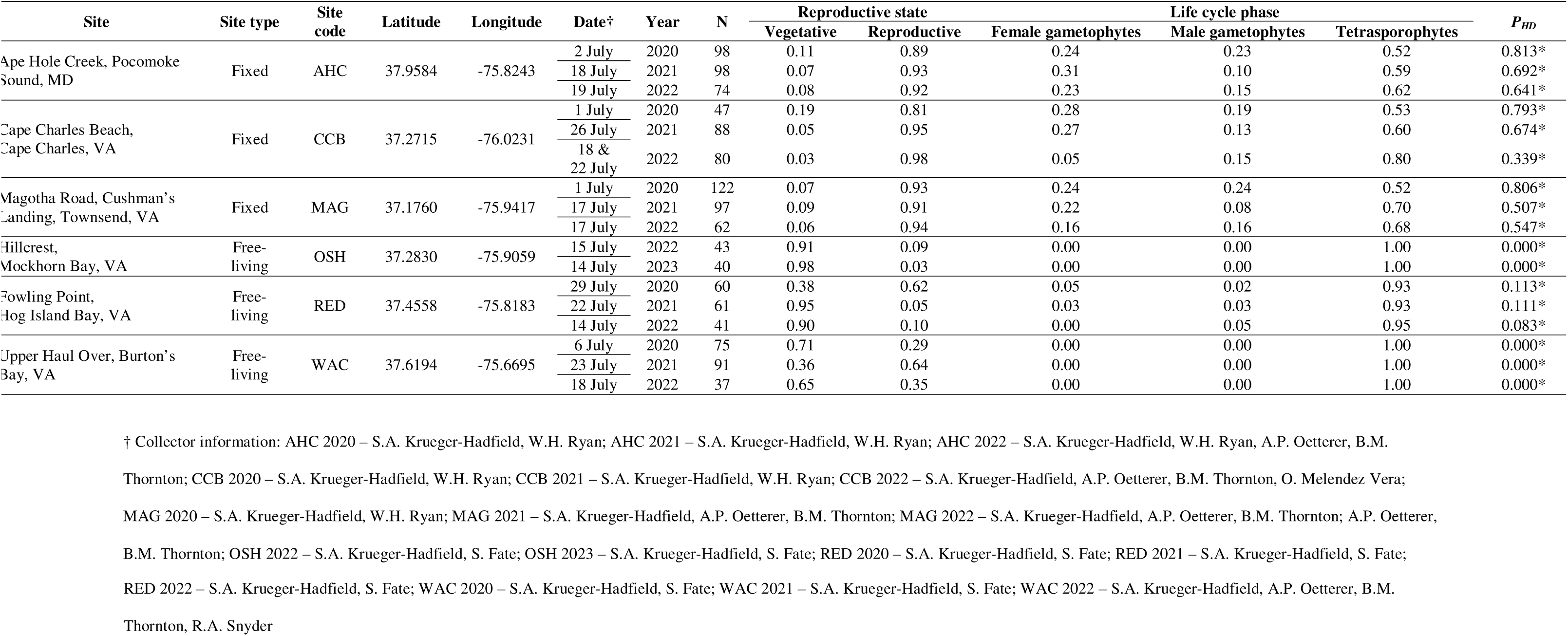
Site information and the proportion of different reproductive states and phases/sexes of *Gracilaria vermiculophylla* at each site and year along the Delmarva Peninsula. Fixed thalli were connected to hard substratum via a holdfast. Free-living thalli drift unattached to substrate or are anchored to a *Diopatra cuprea* tube cap. *P_HD_* is the ploidy diversity metric that describes the amount of phase diversity at each sampling point (Krueger-Hadfield et al., 2019). * denotes phase ratios that were significantly tetrasporophyte biased based on the binomial distribution and deviations from √2:1.

In this study, we explored how hard and soft bottom habitats structured genetic diversity and reproductive modes in populations of *Gracilaria vermiculophylla* along the Delmarva Peninsula, and how these habitats may have shaped and constrained its colonization of this region. We took advantage of a three-year phenology survey from 2020 to 2023 (Krueger-Hadfield et al., 2025b) at sites along the Delmarva Peninsula (Maryland and Virginia, USA) to perform a combination of single time point (i.e., snapshot) and temporal genotyping (i.e., genotyping thalli at two time points) to resolve patterns of reproductive mode variation and genetic structure in *Gracilaria vermiculophylla* (Figure 2). We hypothesized that the invasion of soft sediment habitats results in complex signatures of sexual and clonal reproduction, with concomitant consequences on population structure. In hard bottom habitats, we predicted intergametophytic selfing will prevail due to bottlenecks associated with invasion from Japan and colonization dynamics along the East Coast of the United States. By contrast, soft bottom habitats may be dominated by tetrasporophytes accompanied by signatures of clonal reproduction from thallus fragmentation, though the clonal rate may vary depending on the demographic history of the site. These population genetic signatures will be further magnified by the haploid-diploid life cycle in which the size of the haploid proportion in a population strongly influences patterns of genetic diversity (Stoeckel et al., 2021a). Across all sites, we predict patterns of isolation by distance due to non-motile propagules, though we do expect there to be less differentiation between sites in soft bottom habitats since thalli are not fixed by holdfasts and are likely moving through the interconnected coastal bays of the Delmarva seaside. Within sites in hard bottom habitats, if there are similar allele frequencies between gametophytes and tetrasporophytes (see Engel et al., 2004; Krueger-Hadfield et al., 2011, 2013), then we expect similar levels of genetic differentiation when investigated by pairwise site comparisons separately in the gametophytes or tetrasporophytes. These data will facilitate our understanding of the *G. vermiculophylla* invasion while simultaneously expanding our understanding of reproductive mode variation in haploid-diploid taxa.

## MATERIALS AND METHODS

### Sampling

We collected thalli at three hard bottom sites with fixed thalli (AHC, CCB, and MAG) and two soft bottom sites with free-living thalli (WAC, RED) in July 2020, July 2021, and July 2022 (Table 1; Figure 2). In July 2022 and July 2023, we added an additional soft bottom site with free-living thalli (OSH, Table 1; Figure 2). These collections were part of a wider study of the phenology of fixed and free-living *Gracilaria vermiculophylla* thalli (Krueger-Hadfield et al., 2025b). Approximately 100 thalli were haphazardly collected at hard bottom sites, separated by ∼1 m, and from distinct holdfasts to avoid sampling the same genet twice. Each thallus was individually bagged and kept in spatial sampling order using scroll bags (Krueger-Hadfield, 2021) before processing immediately following collection. At soft bottom sites, thalli were sampled from seven 0.25 m^2^ quadrats. All thalli in each of the quadrats were sampled. In the laboratory, thalli from all sites were carefully disentangled, and a single piece of contiguous thallus was used for downstream analyses.

### Morphological assessment for determining reproductive status and life cycle phase

We determined reproductive state of sampled thalli using a dissecting microscope (40x) to distinguish reproductive (diploid) tetrasporophytes, reproductive male (haploid) gametophytes, and cystocarpic female (haploid) gametophytes (see Krueger-Hadfield et al., 2018). If a thallus had no visible reproductive structures, it was considered vegetative. We preserved a piece of each thallus in silica gel for subsequent DNA extraction. Any cystocarps were excised prior to preservation.

### DNA extraction and PCR amplification

We adapted a Chelex DNA extraction protocol from Simon et al. (2020) as described in Krueger-Hadfield et al. (2023). We placed ∼1 cm of each silica gel-dried thallus into a FisherbrandTM 96-well non-skirted PCR plate and then added 200 µL of pre-heated (95 °C) 5% Chelex solution (Bio-Rad Laboratories, Hercules, CA). Each plate was vortexed for 30 seconds, then incubated at 95 °C for 15 minutes with a brief vortex every 5 minutes. Each plate was centrifuged for 3 minutes at 5,600 x g after which 180 µL of the supernatant was transferred to a new 96-well plate. Each plate was centrifuged again for 3 minutes at 5,600 x g and 150 µL of the supernatant was transferred to a final 96-well plate. For each plate, we had two female gametophytic, two male gametophytic, and two tetrasporophytic positive controls, one negative DNA extraction control, and one negative PCR control.

We used the sex-linked genetic markers developed by Krueger-Hadfield et al. (Krueger-Hadfield et al., 2021b) and modified in Krueger-Hadfield et al. (2023) to determine the phase for each thallus. We amplified the sex-linked loci in a 10 µL duplex PCR containing 2 µL of DNA template, 0.5 U GoTAQ Flexi-DNA Polymerase (Promega, Madison,WI), 1X green reaction buffer, 100 µM of each dNTP, 1.5 mM of MgCl2, 1 mg/mL bovine serum albumin (BSA), 250 nM female forward and reverse primers, and 350 nM male forward and reverse primers, and the cycling profile: 95 °C for 10 minutes, 35 cycles of 95 °C for 30 s, 59 °C for 30 s, and 72 °C for 30 s, followed by a final extension of 72 °C for 5 min. We visualized 3 µL of each PCR product on a 1.5% agarose gel stained with 6 µL GelRed (Biotium, Fremont, CA). We scored each thallus blind and then matched the phase based on the sex-linked marker (female: 73 bp; male: 270 bp; tetrasporophyte: both 73 bp and 270 bp) with the reproductive state determined at the time of collection.

Next, we genotyped the thalli with ten microsatellites described in Kollars et al. (2015) and Krueger-Hadfield et al. (2016). We amplified each locus in simplex in a 10 µL final volume consisting of: 2 µL of DNA template, 0.5 U DreamTaq DNA Polymerase (Promega, Madison, WI), 1X clear reaction buffer, 250 µM of each dNTP, 1.5 mM of MgCl2, 1 mg . mL^-1^ BSA, 100 nM unlabeled forward primer, and 150 nM fluorescently labeled forward primer, 250 nM unlabeled reverse primer, and the cycling profile: 95 °C for 2 minutes, 30 cycles of 95 °C for 30 s, 55 °C for 30 s, and 72 °C for 30 s, followed by a final extension of 72 °C for 5 min (Table S1). For locus Gverm_3003, the concentrations were increased to 350 nM for the unlabeled reverse primer and 250 nM for the fluorescently labeled forward primer. Initial PCR products were diluted in 40 µL UltraPure Distilled Water (Invitrogen, Grand Island, NY, USA). One µL of each PCR product was added to 9 µL of HiDi containing 0.30 µL of size standard (GeneScan500 Liz; Applied Biosystems, Foster City, California, USA) in poolplex as follows: Pool 1: Gverm_1203, Gverm_6311, Gverm_8036, Gverm_804; Pool 2: Gverm_13067, Gverm_5276, Gverm_2790; Pool 3: Gverm_3003, Gverm_1803, Gverm_7969. Fragment analysis was performed at the Heflin Center for Genomic Sciences (hereafter Heflin) at the University of Alabama at Birmingham. Reruns were performed using the same PCR conditions. Fragment analysis was the same as described above except PCR products were not diluted and were electrophoresed on a SeqStudio24 (ThermoFisher Scientific, Waltham, MA) at the Virginia Institute of Marine Science Eastern Shore Laboratory (VIMS ESL).

### Allele calling and genotyping

We ensured that raw allele sizes matched previous genotyping to avoid artificially inflating or deflating allelic diversity and provide a calibration for future studies (see also discussion of calibration in Schoenrock et al. (2020). First, we used thalli that were genotyped by Krueger-Hadfield et al. (2017) and sampled in 2015: five thalli from Hillcrest (OSH), seven thalli from the Upper Haul Over (WAC), and four thalli from Fowling Point (RED). We compared raw and called alleles from Krueger-Hadfield et al. (2017) to those obtained from the Heflin and VIMS ESL platforms and PCRs as described above conducted at VIMS ESL. We note that we used the same fluorochrome for each locus and the same size standard as Krueger-Hadfield et al. (2017). We then used the raw alleles to note any shifts across sequencers. We added thalli from this study sampled at Magotha (MAG) in 2022 (N = 6), Fowling Point in 2020 (N = 6), and the Upper Haul Over in 2021 (N = 4) to ensure that we could compare thalli from the same sites, across sampling years, and across platforms. Loci Gverm_1203, Gverm_6311, Gverm_803, Gverm_13067, and Gverm_5276 were comparable across platforms with no shifts detected (Table S2). Loci Gverm_8036, Gverm_2790, Gverm_3003, and Gverm_7969 all exhibited shifts when comparing Krueger-Hadfield et al. (2017) versus the Helfin and VIMS ESL platforms (Table S2). We removed locus Gverm_1803 (see results) due to problems with accurate allele calling and did not perform a calibration for this locus.

Alleles were manually scored blind using GENEIOUS PRIME v.2023.2.1 (Biomatters, Ltd. Auckland, New Zealand) and we used the allele calls that were calibrated to earlier work as described above (Table S3).

### Phase determination and ratio

As most fragment analysis software assumes diploidy, we considered a thallus as a tetrasporophyte if at least one locus was heterozygous and a thallus as a gametophyte if only one allele was present at all loci (i.e., a fixed homozygote). We then matched the phase designation based on the microsatellite multilocus genotype (MLG) to the reproductive state of the thallus at the time of collection and the phase/sex determined by the sex-linked PCR assay.

The binomial distribution was used to detect deviations from the predicted ratio of √2:1 (0.41 tetrasporophytes to 0.59 gametophytes) for each site (Destombe et al., 1989; C. Thornber & Gaines, 2004). We calculated the ploidy diversity metric (*P_HD_*; Krueger-Hadfield et al., 2019). We used the equation where *x* is the proportion of tetrasporophytes at a site. When *P_HD_* = 1, the ratio of gametophytes to tetrasporophytes is the predicted √2:1, but when *P_HD_* = 0, only gametophytes or sporophytes are present at a site. In our study, as *P_HD_* approaches 0, there are only tetrasporophytes present at a site.

### Genotypic and genetic diversity

We followed the methodological recommendations for analyzing population genetic data in partially clonal taxa as outlined in Krueger-Hadfield & Hoban (2016), Krueger-Hadfield et al. (2021a), and Stoeckel et al. (2021a). We calculated summary statistics at each site and time point using GenAPoPop (Stoeckel et al., 2024a). First, we calculated probability of identity, *pid*, to determine the efficacy of the loci to distinguish between individuals (Krueger-Hadfield et al., 2021a; Stoeckel et al., 2021a; Waits et al., 2001). The *pidu* is the unbiased estimator considering panmixia and *pids* is the estimator for siblings, considering only cross-fertilization between brother and sister. Null allele frequencies were calculated using ML-Null Freq (Kalinowski & Taper, 2006) for the tetrasporophytes and from non-amplification in the gametophytes after discounting technical errors (see Krueger-Hadfield et al., 2011, 2013).

We calculated genotypic richness (*R*) using the formula, where *G* is the number of unique MLGs, and *N* is the total number of thalli genotyped at a site (Dorken & Eckert, 2001) and genotypic evenness (*D*\*) following the equations in Box 3 from Arnaud-Haond et al. (2007). For genotypic evenness, if a site is dominated by a few dominant clones, *D\** approaches 0. In contrast, if at a site, each genet is represented by an equal number of ramets, *D\** will approach 1 even if there are repeated MLGs. To facilitate discussion and interpretation, we explored the relationship between genotypic richness and genotypic evenness following Baums et al. (2006) and Krueger-Hadfield et al. (2021a). We then calculated a multilocus estimate of linkage disequilibrium (, Agapow & Burt, 2001) and the distribution of clonal membership using *Pareto* β (Arnaud-Haond et al., 2007). Krueger-Hadfield et al. (2021a) suggested that *Pareto* β > 2 was associated with low clonal rates, 0.7 < *Pareto* β < 2 was associated with intermediate clonal rates, and *Pareto* β < 0.7 was associated with high clonal rates (see Krueger-Hadfield et al., 2021a, Stoeckel et al. 2021a) Finally, we calculated expected heterozygosity (*H*_E_, Nei, 1978), observed heterozygosity (*H*_O_, Hardy, 2016), and the inbreeding coefficient (*F_IS_*, Weir & Cockerham, 1984) for each locus and the variance associated with each summary statistic (see discussion of this in Stoeckel et al. 2021a). Observed heterozygosities and inbreeding coefficients were only calculated for the tetrasporophytes.

We inferred clonal, selfing, and outcrossing rates at each site using the transition of genotype frequencies over time and the Bayesian method implemented in ClonEstiMate (Becheler et al., 2017) and extended in GenAPoPop (Stoeckel et al., 2024a). We used the uniform joint priors covering the mutation rate u ∈ [0.001, 0.000001, 0.000000001], the clonal rate *c* ∈ [0, 0.1, 0.2, 0.3, 0.4, 0.5, 0.6, 0.7, 0.8, 0.9, 1], and the selfing rate *s* ∈ [0, 0.1, 0.2, 0.3, 0.4, 0.5, 0.6, 0.7, 0.8, 0.9, 1] to cover the full spectrum of all possible quantitative reproductive modes and mutation rates. We considered outcrossing rates as the complement to one out of the clonal and selfing rates (1). We therefore computed the spectrum of discrete posterior probabilities of the distributions of clonal, and selfing rates, and we represented the resulting *a posteriori* distributions in ternary diagrams, one for each site between two successive sampling times. We also reported the credible intervals including 99% of the posterior probability and the maximum posterior probability of the inferred reproductive mode as the joint values of rate of clonality and rate of selfing per sampled site. We note that selfing is possible in *Gracilaria vermiculophylla* if cross-fertilization occurs between two gametophytes that share the same tetrasporophytic parent (i.e., intergametophytic selfing, Klekowski, 1969). We compared genotype frequencies for 2020 to 2021, 2021 to 2022, and 2020 to 2022 for thalli from AHC, CCB, MAG, RED, and WAC and 2022 to 2023 for thalli from OSH (Table 1).

### Ranking sites by clonality and selfing

We used five genetic indices including genotypic richness (*R*), the distribution of clonal membership (*Pareto ß*), the multilocus estimate of linkage disequilibrium (), the inbreeding coefficient and its variance (*F_IS_* and var[*F_IS_*]) to rank the studied sites from the least to the most clonal (Stoeckel et al. 2021a,b). Following Stoeckel et al. (2024b), we computed the sum of the ranks of genetic indices for each population, noted Σ :

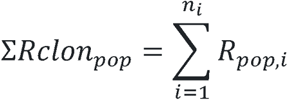

with the number of genetic indices and the rank of the value observed for the population with the set of values for the genetic index, sorted either in an increasing (, var[FIS]) or decreasing (R, Pareto ß, FIS) manner. We also computed the sum of genetic indices normalized between zero and one, noted Σ, which preserves the relative quantitative gap between the studied populations and gives an uniform importance to each index used:

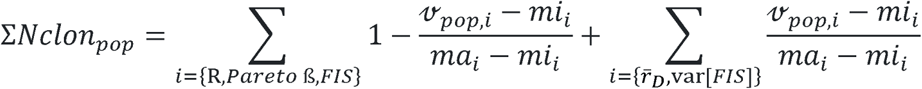

with, the value of the ith genetic index measured in the population, and the minimum and maximum, respectively, value of the ith genetic index measured among the studied populations. Populations with the highest Σ and Σ must be the more clonal.

### Genetic differentiation

To assess genetic divergence between gametophytes and tetrasporophytes (see Engel et al., 2004; Krueger-Hadfield et al., 2011, 2013), we computed average genetic differentiation as *F_ST by phase_* among loci for the two phases in each site in each year following Stoeckel et al. (2021a) and Krueger-Hadfield et al. (2021a). Following Stoeckel et al. 2021b, we also computed for each locus the probability that allele frequencies in gametophytes accurately reflects allele frequencies in tetrasporophytes in a same site !$% as the probability mass function

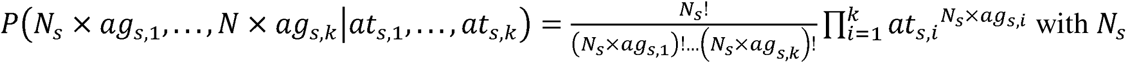

being the number of genotyped gametophytes at the considered site, &, the frequency of allele at site in the genotyped gametophytes, &, the frequency of allele at site in the genotyped tetrasporophytes and k the number of genotyped alleles over all the sampled generations at the considered locus in the considered site. The lower this probability, the more perturbated by migration, by processes subsampling local ancestralities, and potentially by selection at the considered site. We note that this value is a comparison of allele frequencies in the gametophytic and tetrasporophytic subpopulations within a site and not a pairwise comparison between sites as described below.

We estimated pairwise genetic differentiation between the sites by calculating *F*_ST_ (the statistical result from an analysis of molecular variance performed between each pair of sites, Excoffier et al., 1992) and *rho* (an *F_ST_* analogue that is independent of ploidy level, Ronfort et al., 1998) using GenoDive ver. 3.06 (Meirmans, 2020). We performed measures of genetic differentiation between all site types on the tetrasporophytes only since there were few to no gametophytes at free-living sites. We measured geographic distance (km) following the contours of the coastline between all site pairs using the distance function in Google^®^ Maps considering Earth curvature. We then performed Mantel tests to detect relationships between genetic and geographic distance between each site in R.

To further investigate patterns of gene flow between fixed sites and compare between the two phases, we calculated genetic differentiation (*F_ST_*) in the gametophytes using GenoDive.

Finally, the genetic distance between thalli were visualized using a minimum spanning tree of pairwise genetic distances to better interpret genetic ancestry between samples in hierarchically structured populations in the tetrasporophytes between sites using GenAPoPop.

### Comparing summary statistics across sites and phases

We compared summary statistics between fixed and free-living sites and tetrasporophytes and gametophytes at fixed sites using a two-tailed *t*-test and Wilcoxon signed-rank test in R ver. 4.1.3 (R Core Team, 2022) depending on normality and homoscedasticity assumptions to address site- and phase-specific patterns. We also used a Kruskal-Wallis test to identify if the distribution of genetic differentiation per locus varied between and among hard and soft bottom populations.

If there were significant differences in the distributions, for pairwise comparisons, we reported results of Conover’s test of multiple comparisons using rank sums as post hoc test.

### Data visualization

Figures were prepared using R with the following packages: ggplot2 (Wickham, 2016), dplyr (Wickham et al., 2023), reshape2 (Wickham, 2007), ggtext (Wilke & Wiernik, 2022), ggrepel (Slowikowski, 2024), tidyr (Wickham et al., 2024), and matplotlib (Hunter, 2007).

## RESULTS

We genotyped a total of 1,314 thalli from six different sites from 2020 to 2023. Nine thalli (< 1%) did not amplify at a single locus. Thirty-four thalli (< 3%) did not amplify at three or more loci and were removed from the dataset. Another 55 thalli (< 5%) were excluded from the analysis because there was a mismatch between the field identification, sex marker identification, or microsatellite identification that we could not confidently resolve, though this was most likely due to a misidentification of reproductive state. Additionally, the locus Gverm_1803 was removed from the analysis due to inconsistent allele calling and gametophytes with two alleles at this locus. This locus had three alleles all within 2 base pairs (bp) of each other and stuttering that was difficult to score. Of the remaining thalli, 14 tetrasporophytes had missing genotypes at a single locus and one tetrasporophyte had missing genotypes at two loci. These thalli remained in the data set for analyses.

Null alleles were rare based on non-amplification in the gametophytes. Of the 295 gametophytic thalli, one gametophyte did not amplify at locus Gverm_1203, one at locus Gverm_2790, and one at Gverm_3003 after several PCRs. Thus, null allele frequency was less than 1% at these three loci or zero at the other loci. Based on a maximum likelihood estimator in ML-Null Freq, null allele frequency ranged from 0% to 21% in the tetrasporophytes across sites and years (Table S4).

The overall *pidu* for the tetrasporophytes was 3.78 x 10^-6^ and the gametophytes was 4.0 x 10^-5^, and the overall *pids* was 0.02 for the tetrasporophytes and 0.04 for the gametophytes (see Table 2 for site-level estimates of *pidu* and *pids*).

**Table 2a.** Summary statistics based on the *Gracilaria vermiculophylla* tetrasporophyte data set at each site and year. *N*, number of genotyped tetrasporophytes; *pidu*, unbiased probability of identity under panmixia; *pids*, probability of identity between sibs; *G*, number of unique multilocus genotypes (MLGs); *R*, genotypic richness; *D\**, genotypic evenness; *Pareto* β, distribution of clonal membership;, multilocus estimate of linkage disequilibrium; *H*_E_ mean multilocus expected heterozygosity and its variance across loci shown in parentheses; *H*_O_, mean multilocus observed heterozygosity and its variance shown in parentheses; and *F_IS_*, mean multilocus inbreeding coefficient and its variance across loci shown in parentheses.

| Site | Year | $N$ | $pidu$ | $pids$ | $G$ | $R$ | $D^*$ | *Pareto* $\beta$ | $\bar{r}_D$ | $H_E$ (var) | $H_O$ (var) | $F_{IS}$ (var) |
| --- | --- | --- | --- | --- | --- | --- | --- | --- | --- | --- | --- | --- |
| AHC | 2020 | 51 | $9.68 \times 10^{-3}$ | 0.184 | 31 | 0.600 | 0.963 | 0.916 | 0.081 | 0.179 (0.285) | 0.176 (0.308) | 0.012 (2.067) |
| | 2021 | 58 | $2.36 \times 10^{-2}$ | 0.261 | 24 | 0.404 | 0.930 | 0.642 | 0.072 | 0.142 (0.232) | 0.132 (0.220) | 0.069 (3.535) |
| | 2022 | 46 | $1.45 \times 10^{-3}$ | 0.121 | 32 | 0.689 | 0.980 | 1.560 | 0.127 | 0.214 (0.419) | 0.181 (0.280) | 0.154 (1.656) |
| CCB | 2020 | 25 | $1.06 \times 10^{-4}$ | 0.037 | 24 | 0.958 | 0.997 | 3.644 | 0.090 | 0.335 (0.526) | 0.311 (0.518) | 0.070 (1.940) |
| | 2021 | 53 | $9.36 \times 10^{-5}$ | 0.032 | 52 | 0.981 | 0.999 | 4.728 | 0.047 | 0.348 (0.570) | 0.358 (0.625) | -0.030 (1.092) |
| | 2022 | 64 | $5.50 \times 10^{-6}$ | 0.021 | 63 | 0.984 | 1.000 | 5.000 | 0.044 | 0.376 (0.698) | 0.345 (0.644) | 0.082 (1.251) |
| MAG | 2020 | 64 | $4.81 \times 10^{-5}$ | 0.035 | 63 | 0.984 | 1.000 | 5.000 | 0.072 | 0.338 (0.573) | 0.333 (0.533) | 0.016 (0.986) |
| | 2021 | 68 | $2.14 \times 10^{-5}$ | 0.031 | 62 | 0.910 | 0.996 | 2.270 | 0.030 | 0.346 (0.584) | 0.356 (0.639) | -0.030 (1.163) |
| | 2022 | 42 | $2.50 \times 10^{-6}$ | 0.018 | 42 | 1.000 | 1.000 | 4.426 | 0.046 | 0.393 (0.682) | 0.344 (0.541) | 0.124 (0.900) |
| OSH | 2022 | 43 | $3.06 \times 10^{-6}$ | 0.023 | 29 | 0.667 | 0.973 | 0.925 | 0.183 | 0.376 (0.396) | 0.352 (0.416) | 0.064 (1.212) |
| | 2023 | 40 | $2.83 \times 10^{-7}$ | 0.023 | 21 | 0.513 | 0.937 | 0.780 | 0.269 | 0.375 (0.445) | 0.400 (0.543) | -0.067 (1.222) |
| RED | 2020 | 56 | $6.61 \times 10^{-5}$ | 0.046 | 38 | 0.673 | 0.958 | 0.655 | 0.168 | 0.309 (0.516) | 0.319 (0.635) | -0.034 (0.220) |
| | 2021 | 57 | $-5.20 \times 10^{-9}$ | 0.007 | 48 | 0.839 | 0.990 | 1.526 | 0.203 | 0.475 (0.673) | 0.495 (0.752) | -0.043 (0.152) |
| | 2022 | 39 | $8.77 \times 10^{-7}$ | 0.010 | 20 | 0.500 | 0.900 | 0.497 | 0.356 | 0.453 (0.517) | 0.544 (1.124) | -0.202 (1.717) |
| WAC | 2020 | 75 | $2.02 \times 10^{-3}$ | 0.116 | 23 | 0.297 | 0.779 | 0.219 | 0.189 | 0.220 (0.464) | 0.273 (0.982) | -0.237 (1.297) |
| | 2021 | 91 | $2.05 \times 10^{-5}$ | 0.046 | 53 | 0.578 | 0.914 | 0.354 | 0.146 | 0.305 (0.554) | 0.291 (0.910) | 0.048 (1.634) |
| | 2022 | 37 | $2.96 \times 10^{-3}$ | 0.117 | 10 | 0.250 | 0.578 | 0.124 | 0.524 | 0.222 (0.412) | 0.270 (1.126) | -0.219 (3.159) |

**Table 2b.** Summary statistics based on the *Gracilaria vermiculophylla* gametophyte data set at each fixed site and year. *N*, number of genotyped gametophytes; *pidu*, unbiased probability of identity under panmixia; *pids*, probability of identity between sibs; *G*, number of unique multilocus genotypes (MLGs); *R*, genotypic richness; *D\**, genotypic evenness; *Pareto* β, distribution of clonal membership; and multilocus *H*_E_, expected heterozygosity and variance shown in parentheses.

| Site | Year | *N* | *pidu* | *pids* | *G* | *R* | *D\** | *Pareto β* | *H<sub>E</sub>* (var) |
| --- | --- | --- | --- | --- | --- | --- | --- | --- | --- |
| AHC | 2020 | 47 | 5.65 x 10<sup>-3</sup> | 0.171 | 16 | 0.326 | 0.845 | 0.373 | 0.185 (0.280) |
|  | 2021 | 40 | 6.13 x 10<sup>-3</sup> | 0.169 | 16 | 0.385 | 0.860 | 0.411 | 0.188 (0.240) |
|  | 2022 | 28 | 1.38 x 10<sup>-3</sup> | 0.114 | 14 | 0.481 | 0.907 | 0.667 | 0.221 (0.393) |
| CCB | 2020 | 22 | 9.64 x 10<sup>-6</sup> | 0.039 | 19 | 0.857 | 0.983 | 1.848 | 0.323 (0.570) |
|  | 2021 | 35 | 7.61 x 10<sup>-5</sup> | 0.046 | 27 | 0.765 | 0.985 | 2.127 | 0.307 (0.625) |
|  | 2022 | 16 | 7.27 x 10<sup>-6</sup> | 0.022 | 15 | 0.933 | 0.992 | 3.000 | 0.369 (0.700) |
| MAG | 2020 | 58 | 2.30 x 10<sup>-5</sup> | 0.027 | 45 | 0.772 | 0.989 | 1.862 | 0.358 (0.642) |
|  | 2021 | 29 | 3.45 x 10<sup>-5</sup> | 0.031 | 26 | 0.893 | 0.993 | 2.273 | 0.355 (0.616) |
|  | 2022 | 20 | 5.32 x 10<sup>-5</sup> | 0.029 | 18 | 0.895 | 0.989 | 2.322 | 0.357 (0.641) |

### Phase ratio and ploidy diversity

Across all sites and years, reproductive thalli ranged from 3% to 98% in the sample from each population and time point (Reproductive state, Table 1; see Krueger-Hadfield et al., 2025b). All sites and time points were significantly tetrasporophyte-biased. Gametophytes were nevertheless common at hard bottom sites, accounting for anywhere from 20% to 48% of the sampled thalli (Life cycle phase, Table 1). However, at the three soft bottom sites, OSH, RED, and WAC, we found tetrasporophytes accounted for 93% or greater of the thalli sampled (Table 1). We only found gametophytes at low frequency (< 7%) at RED as drifting thalli or anchored to *Diopatra cuprea* tube caps (see definitions in Krueger-Hadfield et al., 2023, 2025a). There was a significant difference between hard and soft bottom sites as assessed by *P_HD_* (W = 72, *p* < 0.001).

### Genotypic diversity

Across all sites and time points, there were 513 unique multilocus genotypes (MLGs) out of 909 tetrasporophytes genotyped. We re-encountered 93 MLGS anywhere from once to 89 times within and across sites and time points (Table S5a). The majority of these MLGs would be considered clones based on *p_sex_*whereby 74 (∼80%) had *p*-values < 0.05, 8 (∼8%) had *p*-values between 0.05 and 0.09, and 11 (∼12%) had *p*-values > 0.1 (Table S5a). Sixteen MLGs were found between two and seven times at the same site and same year and 30 MLGs were sampled at the same site but over different sampling years (21 in two years and 9 across all three years). All other repeated MLGs were re-encountered at more than one site, with 16 MLGs found at two sites and two years, 12 at two sites and three years or four years (OSH was only sampled in 2022 and 2023), and eight sampled at three sites and in different sampling years. Nine of the MLGs with significant *p_sex_* values were found at both hard and soft bottom sites, including at CCB and AHC (Table S5a).

For the fixed tetrasporophytes, genotypic richness (*R*) for the tetrasporophytes at hard bottom sites was between 0.40 to 1.0 (Table 2a, Figure 3a). CCB and MAG had more unique tetrasporophytes as compared to AHC, where there were many more repeated genotypes. Most repeated MLGs at AHC were sampled throughout the sampling area, although there were several examples in which repeated MLGs were sampled in consecutive order (e.g., MLG137, see Table S5a). Notably, the *p_sex_*value for the re-encountered genotypes for this MLG was not < 0.05. For the free-living tetrasporophytes, the genotypic richness ranged from 0.25 to 0.94, with the thalli at WAC more often sharing the same genotype. Repeated MLGs at WAC were more often found within the same quadrat than RED or OSH. We re-encountered some MLGs between hard and soft bottom sites including within and between years (Table S5a). Overall, fixed tetrasporophytes had higher genotypic richness as compared to free-living tetrasporophytes (Figure 3a; W = 62, p = 0.014).

**Figure 3.**
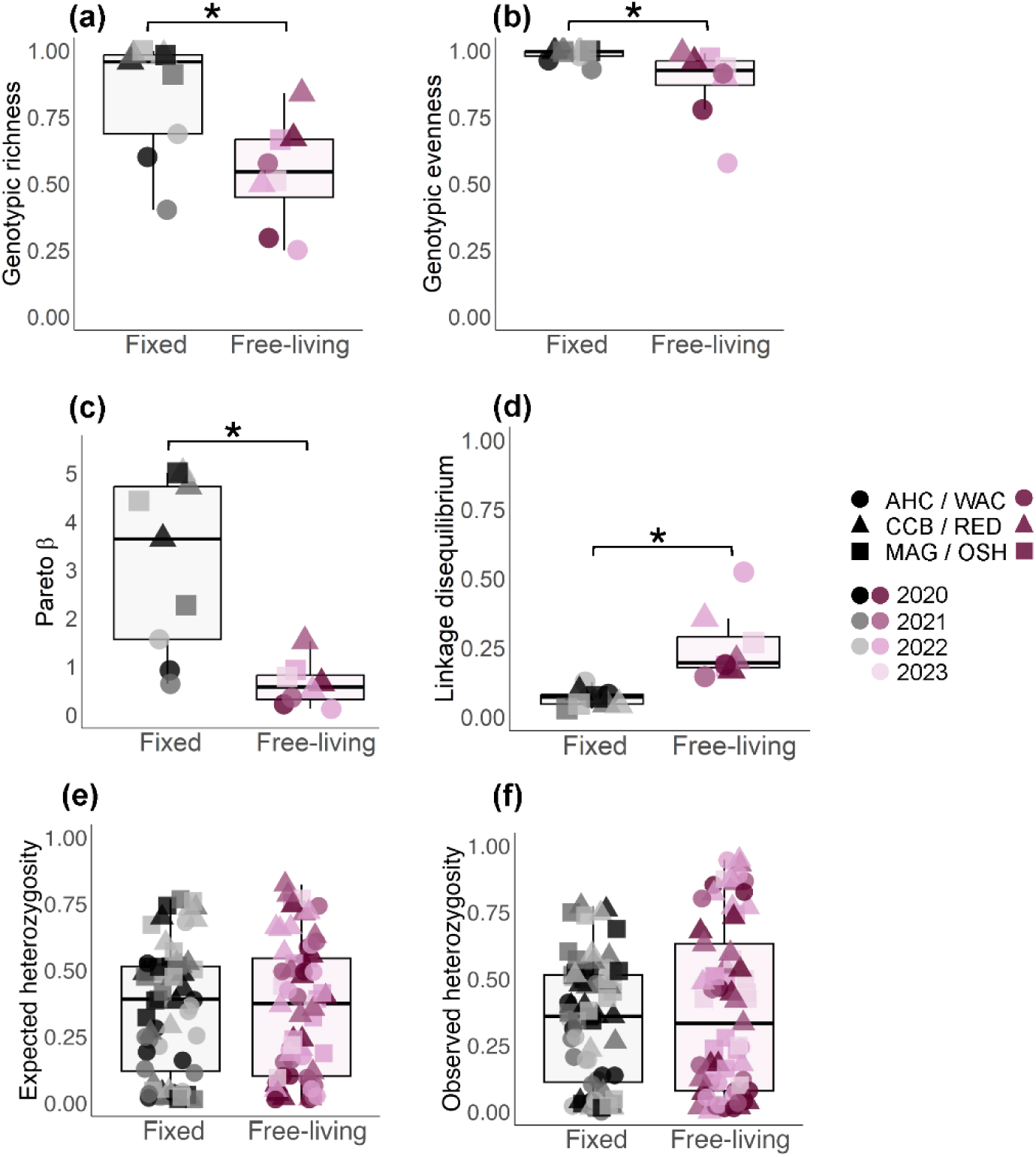
Boxplots of single locus values of (a) genotypic richness (*R*), (b) genotypic evenness (*D\**), (c) expected heterozygosity (*H_E_*), (d) observed heterozygosity (*H_O_*), (e) the distribution of clonal membership (*Pareto ß*), and (f) the multilocus estimate of linkage disequilibrium () depending if sampled populations were found fixed (black colors) or free-living (maroon colors) in their habitats. Expected and observed heterozygosity are shown as singe locus values. Sites are represented as shapes and colors indicate sampling year, with hard bottom sites with fixed thalli shown in shades of black and soft bottom sites with free-living thalli shown in shades of maroon. Boxes represent the interquartile range with the line inside the box indicating the median.

Across all sites and time points, there were 121 unique multilocus genotypes (MLGs) out of 295 gametophytes genotyped (Table S5b). We re-encountered 29 MLGs anywhere from once to 42 times within and across sites and time points (Table S5b). Unlike the tetrasporophytes, the majority of these MLGs would not be considered clones based on *p_sex_* whereby 11 (∼38%) were < 0.05, 1 (∼3%) was between 0.05 and 0.09, and 17 (∼59%) were > 0.1 (Table S5B). Of the repeated MLGs with *p*-values < 0.05, two were re-encountered once, one of which was found at MAG in 2020 and 2021. The other was found at CCB in 2021 and MAG in 2022. All other re-encountered MLGs that had significant *p*-values were found across two or three sites and different years. Repeated MLGs sampled in the same year and same site were mostly sampled across the sampling site, although MLG012 (21 thalli), MLG019 (21 thalli), and MLG028 (43 thalli) have some re-encounters in close numerical order.

Gametophytic genotypic richness (*R*) ranged from 0.33 to 0.93 (Table 2b, Figure 4a). CCB and MAG had higher genotypic richness as compared to AHC. There was no significant difference between tetrasporophytes and gametophytes in genotypic richness (Figure 4a; W = 61, *p* = 0.077), though mean genotypic richness tended to be higher in tetrasporophytes.

**Figure 4.**
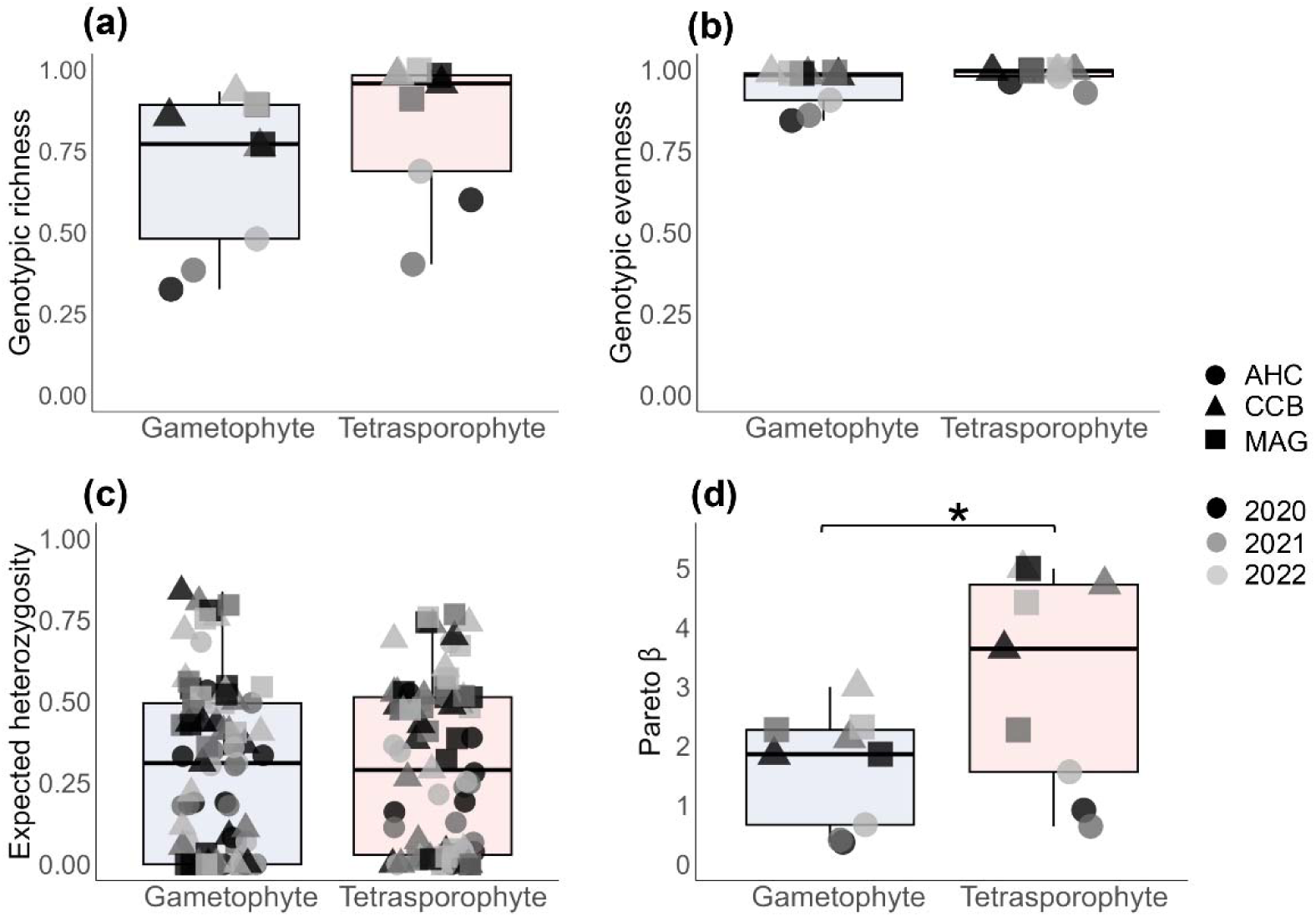
Boxplots of single locus values of (a) genotypic richness (*R*), (b) genotypic evenness (*D\**), (c) expected heterozygosity (*H_E_*), and (d) the distribution of clonal membership (*Pareto ß*), and (E) the multilocus estimate of linkage disequilibrium () are shown for gametophytes (blue) and tetrasporophytes (red). Expected heterozygosity are shown as singe locus values. Sites are represented as shapes and colors indicate sampling year. Boxes represent the interquartile range with the line inside the box indicating the median.

Genotypic evenness (D*) was high across all sites for gametophytes and tetrasporophytes (0.845 < D* < 1, Table 2, Figure 3b, Figure 4b), except for WAC tetrasporophytes in 2020 and 2022 (Table 2a). Genotypic evenness was lower in soft bottom sites (i.e., there were some genets that dominated soft bottom sites; W = 65, *p* = 0.006) and almost significantly lower in gametophytes as compared to tetrasporophytes (W = 63, *p* = 0.052).

Unlike previous comparisons of non-native sites, there were no clear groupings of sites into more sexual (i.e., hard bottom sites with fixed thalli) versus more clonal (i.e., soft bottom sites with free-living thalli) when comparing the relationship between genotypic richness and evenness (Figure S1). Tetrasporophytic thalli at CCB and MAG were clustered in the top right of the plot with high genotypic richness and evenness, indicative of sexual reproduction. By contrast, gametophytes and AHC thalli were characterized by variable genotypic richness, but generally high evenness.

*Pareto* β was greater in hard bottom sites as compared to soft bottom sites for the tetrasporophytes (Table 2a, Figure 3c; *t_9.13_* = 4.03, *p* < 0.003). The *Pareto ß* values for soft bottom sites ranged from 0.124 to 1.526 for tetrasporophytes. *Pareto ß* was greater for tetrasporophytes than gametophytes (Figure 4c; *t_12.10_* = 2.17, *p* = 0.049), where the *Pareto ß* range was from 0.642 to 5.000 for tetrasporophytes (Table 2a) and 0.373 to 3.000 for gametophytes (Table 2b). AHC was characterized by *Pareto ß* values of < 2, whereas CCB and MAG had values close to or greater than 2.

Linkage disequilibrium () was greater in free-living (0.146 to 0.524) than in fixed tetrasporophytes (0.03 to 0.127; Table 2b, Figure 3d; W = 0; *p* < 0.001).

### Genetic diversity

Expected heterozygosity (*H_E_*; W = 2292, *p* = 0.934) was similar between fixed and free-living tetrasporophytes (Figure 3e, Table 2, Table S6a). Observed heterozygosity (*H_O_*; W = 2167.5, *p* = 0.532) was almost significantly greater in free-living tetrasporophytes as compared to fixed (Figure 3e, Table 2, Table S6b). However, AHC thalli were consistently represented by low *H_E_* and *H_O_* values (Table 2, Table S6a) as compared to the other sites. There was no difference in expected heterozygosity between gametophytes and tetrasporophytes (Figure 4d, Table 2, Table S6a; W = 3402, *p* = 0.684).

Multilocus mean *F_IS_* values were slightly positive in fixed sites and slightly negative in free-living sites (Table 2a, Table S6c), though there was no difference between site types (W = 2727, *p* = 0.101). Single locus values varied across sites (Figure 5; -0.261 ≤ fixed tetrasporophytic *F_IS_* < 1.000; -0.805 ≤ free-living tetrasporophytic *F_IS_* < 1.000). The *F_IS_* variance ranged from 0.900 to 3.535 (mean = 1.621) at hard bottom sites and from 0.152 to 3.159 (mean = 1.327) at soft bottom sites. There was no difference between the variance in *F_IS_* in fixed and free-living tetrasporophytes (*W* = 40, *p* = 0.736).

**Figure 5.**
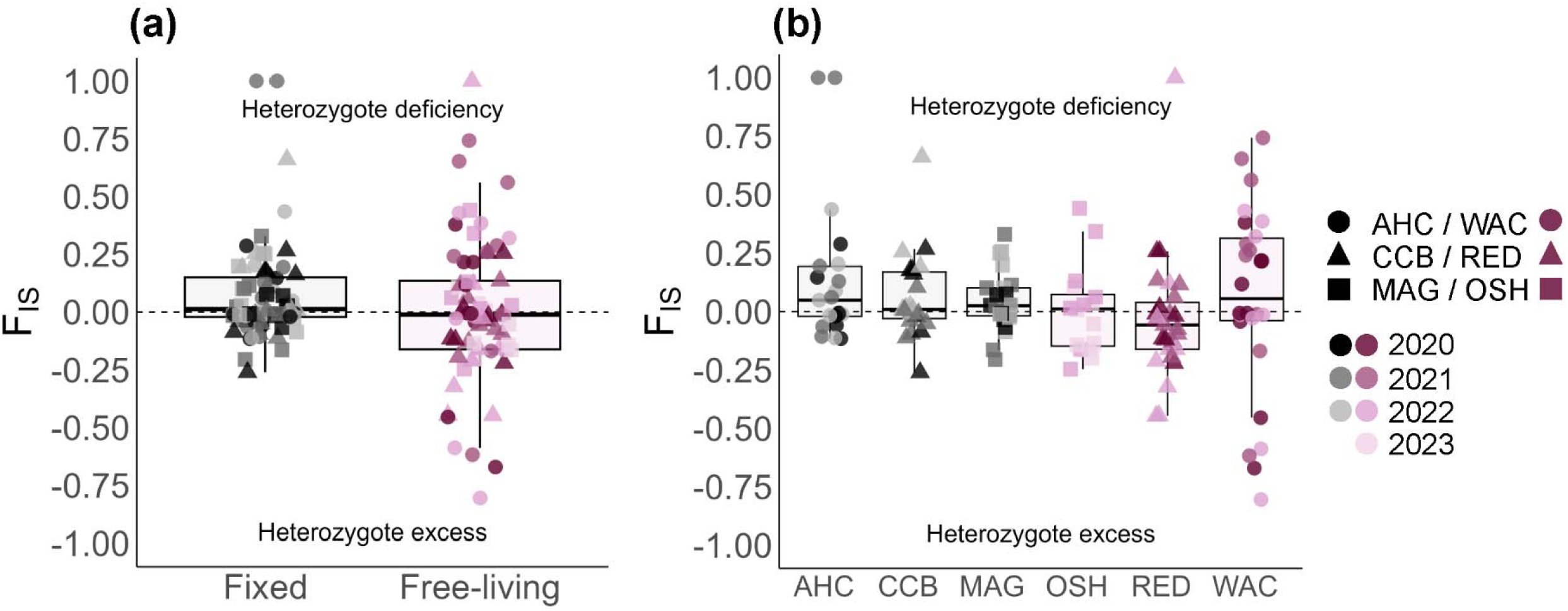
Single locus values of the inbreeding coefficient (*F*_IS_) in (a) in fixed thalli in hard bottom sites and free-living thalli in soft bottom sites and (b) for each site. Sites are indicated as shapes and shades of colors indicate sampling year. Populations attached to hard substrates are shown in shades of black, while those that were free-living within their habitats are represented in shades of maroon. The dashed line at *F_IS_* = 0 shows expected Hardy–Weinberg proportions homozygotes and heterozygotes. Boxes represent the interquartile range with the line inside the box indicating the median.

Overall, genetic differentiation (*F_ST_*) between phases were low (5.66 x 10^-5^ ≤ *F_ST by phase_* ≤ 6.68 x 10^-4^) and of similar magnitude regardless of site or year (Table 3). Locally, gametophytes and tetrasporophytes are linked, but there was variation by locus. Some loci at a given site and time point, such as locus Gverm_2790 at MAG 2020, showed expected local probability of transition of allele frequency between gametophytes and tetrasporophytes (=1). Other loci, such as Gverm_13067 at CCB 2022, showed low probability of transition of allele frequency between gametophytes and tetrasporophytes (=3.91 x 10^-31^), indicating a possible disjunction of allele frequencies between the phases.

**Table 3.** Genetic differentiation between gametophytes and tetrasporophytes as measured by *F_ST_*, genetic distance, and the probability by locus that allele frequencies in gametophytes were drawn from allele frequencies in tetrasporophytes (Multinomial pmf).

| Site | Year | $F_{ST}$ | Genetic<br>differentiation | Multinomial pmf | | | | | | | | |
| --- | --- | --- | --- | --- | --- | --- | --- | --- | --- | --- | --- | --- |
|  |  |  |  | Gverm_1203 | Gverm_6311 | Gverm_8036 | Gverm_804 | Gverm_13067 | Gverm_5276 | Gverm_2790 | Gverm_3003 | Gverm_7969 |
| AHC | 2020 | $5.66 \times 10^{-5}$ | 0.0165 | 0.1695 | 0.1002 | 0 | 0.1855 | 1.0 | $8.01 \times 10^{-76}$ | 0.6294 | 0 | 1.0 |
| AHC | 2021 | $1.30 \times 10^{-4}$ | 0.0307 | 0.1289 | 0.0496 | 0.0248 | 0.1578 | $2.90 \times 10^{-71}$ | $1.38 \times 10^{-65}$ | 1.0 | $5.94 \times 10^{-64}$ | 1.0 |
| AHC | 2022 | $1.57 \times 10^{-4}$ | 0.0214 | $6.03 \times 10^{-39}$ | $3.60 \times 10^{-48}$ | $1.51 \times 10^{-12}$ | $1.58 \times 10^{-43}$ | 0.2266 | $8.36 \times 10^{-5}$ | 1.0 | 0.1681 | 1.0 |
| CCB | 2020 | $3.84 \times 10^{-4}$ | 0.0374 | $8.05 \times 10^{-18}$ | 0.0412 | 0.0670 | $8.05 \times 10^{-18}$ | 0.2879 | 0 | 1.0 | 0.0179 | 1.0 |
| CCB | 2021 | $6.68 \times 10^{-4}$ | 0.0471 | $4.27 \times 10^{-34}$ | 0.0308 | 0.0368 | $2.03 \times 10^{-38}$ | 0.3571 | 0 | 0.7177 | 0.0057 | 1.0 |
| CCB | 2022 | $3.08 \times 10^{-4}$ | 0.0450 | 0.0154 | $8.79 \times 10^{-18}$ | $1.44 \times 10^{-51}$ | $1.33 \times 10^{-17}$ | $3.91 \times 10^{-31}$ | $4.74 \times 10^{-16}$ | 1.0 | 0.0483 | $1.93 \times 10^{-34}$ |
| MAG | 2020 | $2.02 \times 10^{-4}$ | 0.0250 | 0.0032 | $1.21 \times 10^{-37}$ | 0.0124 | 0.0089 | 0.3693 | $2.15 \times 10^{-22}$ | 1.0 | 0.0006 | 0.6345 |
| MAG | 2021 | $2.74 \times 10^{-4}$ | 0.0328 | $1.75 \times 10^{-17}$ | 0.0764 | $1.97 \times 10^{-15}$ | $7.44 \times 10^{-19}$ | 0.2817 | $2.66 \times 10^{-16}$ | 0.8073 | 0.0004 | 1.0 |
| MAG | 2022 | $1.57 \times 10^{-4}$ | 0.0527 | $6.40 \times 10^{-12}$ | $1.23 \times 10^{-10}$ | $5.92 \times 10^{-15}$ | $2.47 \times 10^{-20}$ | 0.6176 | $6.23 \times 10^{-6}$ | 1.0 | $1.50 \times 10^{-5}$ | $3.27 \times 10^{-39}$ |

### Temporal analyses of the reproductive mode

Inferred clonal rates were greater in the three soft bottom sites with free-living tetrasporophytes as compared to the hard bottom sites with fixed tetrasporophytes (_. /_=0.5, _. 0 1 %_ = 29.5, *p* = 0.009, medians: _. /_ = 0.09, _. 0 1 %_ = 0.84; Table 4, Figure 6, Table S7). We found no difference in inferred selfing rates between fixed and free-living tetrasporophytes (= 21, _. 0 1 %_ = 9, *p* =0.329, medians:= 0.39, _. 0 1 %_ = 0.14). Fixed thalli in hard bottom sites had signatures of greater clonality than selfing (*W* = 0, *p* = 0.031). Between 2020 – 2021 and 2021 – 2022, an increase in clonality was marginally non-significant (*W* = 0, *p* = 0.062) while selfing rates did not change (*W* = 4, *p* = 0.219).

**Figure 6.**
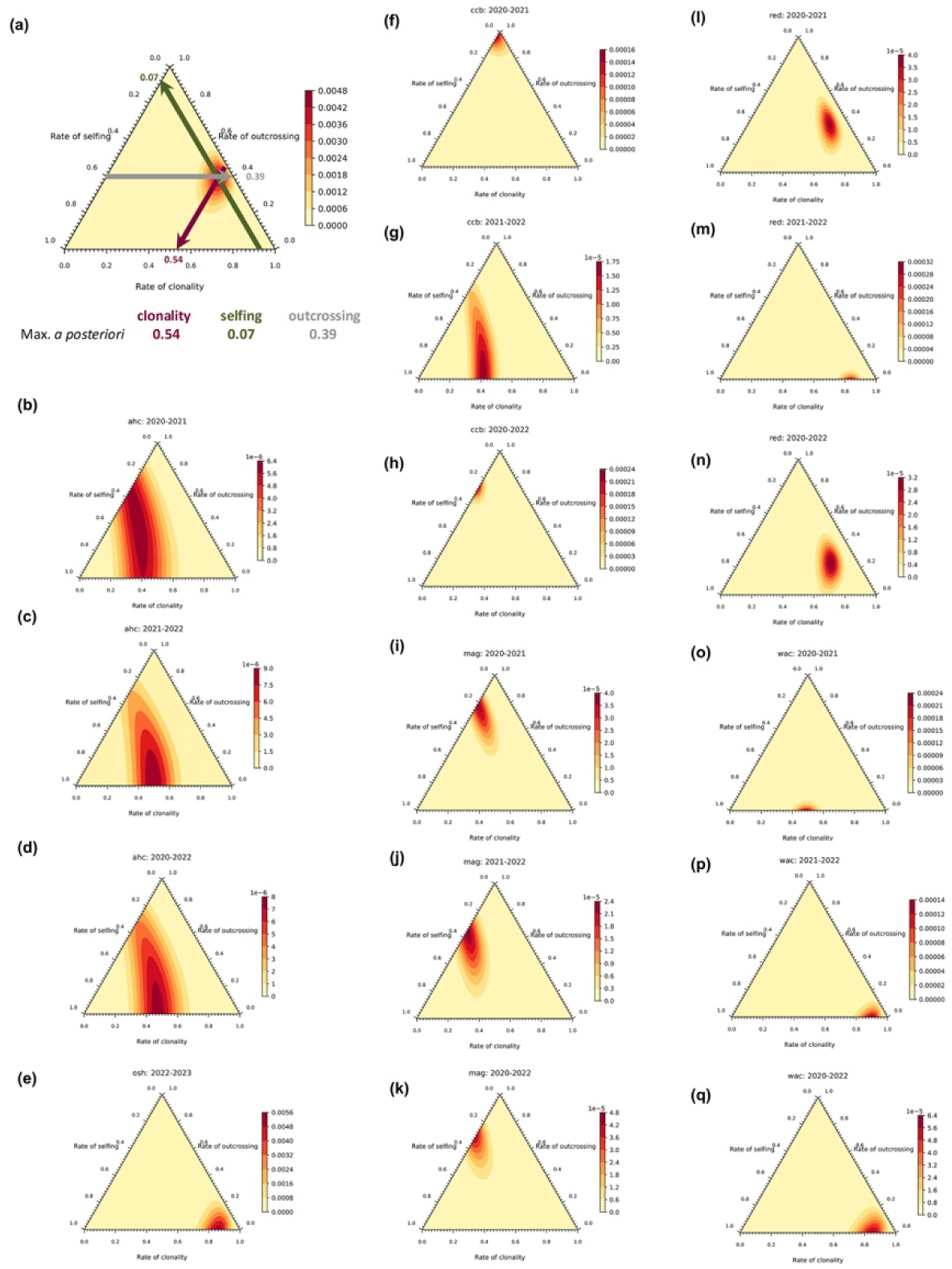
Ternary heat-map plots of the posterior probabilities of rates of selfing, outcrossing, and clonality using the change of genotype frequencies between temporal samples for each site and sampling interval combination. (a) an example ternary plot showing the clines in selfing, outcrossing, and clonality. The clines are drawn parallel to the sides of the triangle such that they cross the maximum posterior probability (i.e., the red center of the heat map) to obtain the value for the considered axis (i.e., rate of selfing, outcrossing, or clonality). Ternary plots by site and temporal comparison: (b) Ape Hole Creek (AHC) 2020 – 2021, (c) AHC 2021 – 2022, (d) AHC 2020 – 2022, (e) OSH 2022-2023, (f) Cape Charles Beach (CCB) 2020 – 2021, (g) CCB 2021 – 2022, (h) CCB 2020 2022, (i) Magotha Road (MAG) 2020 – 2021, (j) MAG 2021 – 2022, (k) MAG 2020 – 2022, (l) Fowling Point (RED) 2020 – 2021, (m) RED 2021 – 2022, (n) RED 2020 – 2022, (o) Upper Haul Over (WAC) 2020 – 2021, (p) WAC 2021 – 2022, and (q) WAC 2020 –2022.

**Table 4.**
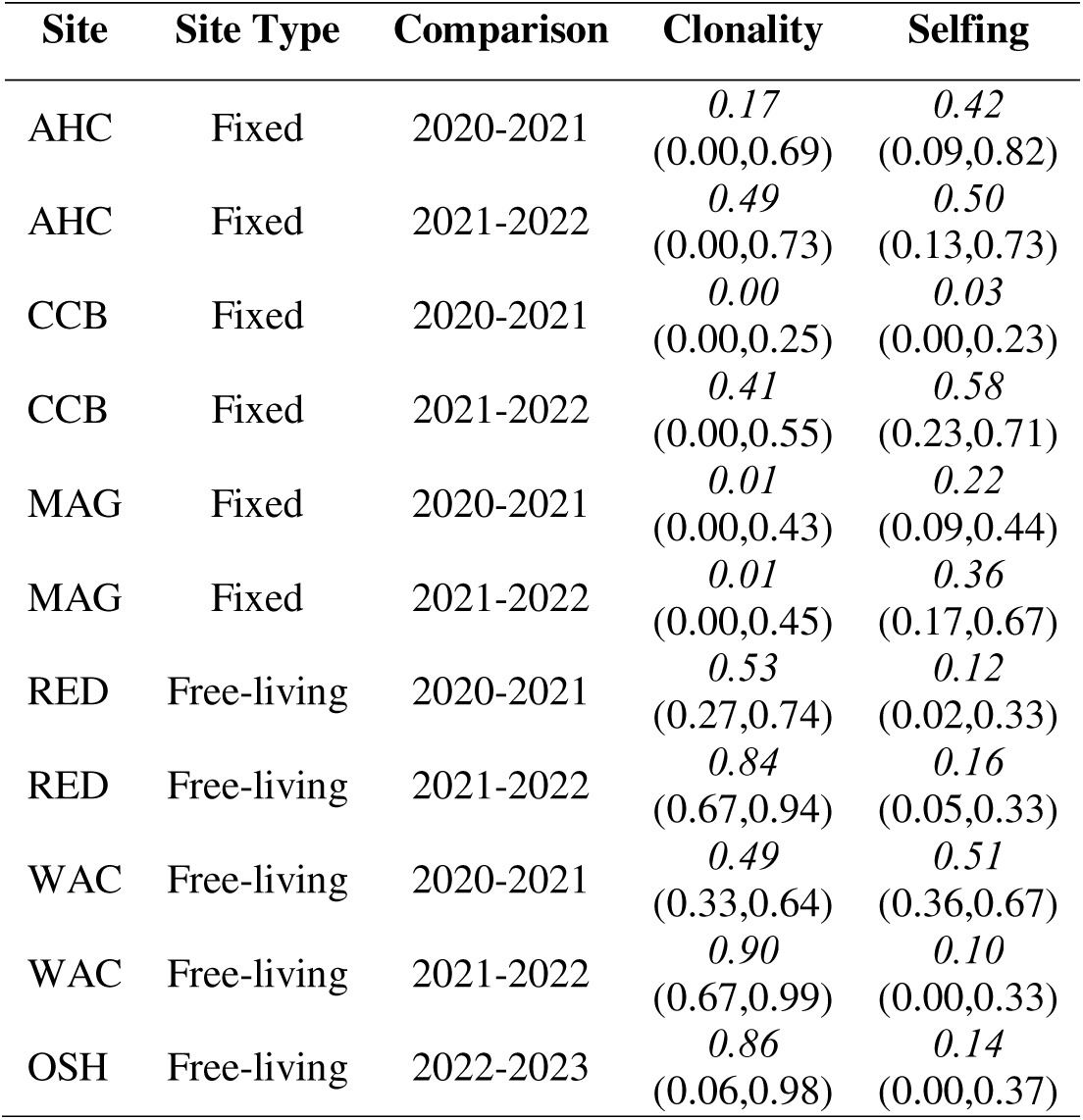
Maximum posterior probabilities (in italic) and credible intervals including more than 99% of the posterior probabilities (in parenthesis) of rates of clonality and rates of selfing inferred in each site using transitions of genotype frequencies between two successive temporal samples.

Outcrossing rates declined over time at MAG, though outcrossing rates were greater at MAG than at AHC. AHC also was characterized by the largest credible interval and in the 2020 – 2021 comparison, three differing rates of selfing and outcrossing. The clonal and selfing rates may be obscured by high selfing rates at AHC. At CCB, we observed a switch from outcrossing to selfing from 2020 – 2021 and 2021 – 2022. At RED, there was an increasing clonal rate over time. In the 2020 – 2022 comparison, we would have expected to see an increasing rate of sexuality, but at RED, we observed an increasing rate of clonality and selfing, suggesting selfing was indeed occurring. Two generations of selfing (i.e., two years between sampling time points) confuses clonal and selfing rates. In contrast, the clonal rates increased at WAC over time, and selfing rates declined.

### Ranking sites by clonality and selfing

We were able to disentangle three groups of sites that were stable across the sampling time points based on genotypic richness (*R*), *Pareto ß*, linkage disequilibrium (), the inbreeding coefficient (*F_IS_*), and the variance in the inbreeding coefficient (var[*F_IS_*]; Table S8). The hard bottom sites MAG and CCB were in the group that was more sexual with a significant but low level of clonality. These two sites had high genotypic diversity, high values of *pareto ß*, low linkage disequilibrium, zero or slightly positive *F_IS_* values, and low *F_IS_* variance. AHC, RED, and OSH made up the group with intermediate clonal rates. For RED and OSH, both sites with free-living tetrasporophytes, low genotypic diversity, low values of *pareto ß*, significant linkage disequilibrium, and high *F_IS_* variance, coupled with tetrasporophytic bias all point to clonality.

However, AHC is a hard bottom site with fixed thalli, suggesting the low genotypic diversity, low values of *pareto ß*, significant linkage disequilibrium, and high *F_IS_* variance may be more indicative of self-fertilization against a backdrop of a clonal legacy on the genome. WAC had the highest clonal ranking and is also the most stable free-living site in terms of *Gracilaria vermiculophylla* biomass (Krueger-Hadfield et al., 2025b).

### Genetic structure

Within sampling year, there were no significant differences between estimates of genetic differentiation as measured by *F_ST_* in the tetrasporophytes when comparing between fixed tetrasporophytes, free-living tetrasporophytes, or fixed versus free-living tetrasporophytes (*F_2,32_* = 1.93, *p* = 0.162). There were, however, some patterns suggesting a trend of lower genetic differentiation between soft bottom sites in 2020 and 2021, though the amount of differentiation was almost twice as much when comparing allele frequencies at RED and WAC in 2022 (Table 5a, Table S9a-h). Nevertheless, genetic differentiation was generally twofold greater in the pairwise comparisons between hard bottom sites and between hard and soft bottom site pairwise comparisons as compared to those between soft bottom sites (Table 5a, Table S9a-h). As geographic distance increased, so did genetic distance as measured by *F_ST_* (Figure 7) and *rho* (Figure S2), regardless of year.

**Table 5a.** Pairwise *F_ST_*means and standard error of site type comparisons for 2020-2022/2023 and tetrasporophytes. Numbers in parentheses are the number of pairwise comparisons. There were only two free-living sites sampled in 2020 and 2021, resulting in one pairwise comparison for those years. Fixed sites (AHC, CCB, MAG) are characterized by thalli fixed to hard substratum via a holdfast. Free-living sites (OSH, RED, WAC) are characterized by thalli drifting unattached to substrate, or anchored to a *Diopatra cuprea* tube cap.

| 1337 | Site comparisons | 2020 | 2021 | 2022 | All years |
| --- | --- | --- | --- | --- | --- |
|  | Fixed vs. Fixed | 0.133 ± 0.061 (3) | 0.151 ± 0.068 (3) | 0.098 ± 0.049 (3) | 0.127 ± 0.031 (9) |
| 1338 | Free-living vs. Free-living | 0.056 (1) | 0.085 (1) | 0.144 ± 0.031 (3) | 0.115 ± 0.025 (5) |
|  | Fixed vs. Free-living | 0.181 ± 0.038 (6) | 0.182 ± 0.028 (6) | 0.175 ± 0.032 (9) | 0.179 ± 0.018 (21) |
| 1339 | Overall | 0.154 ± 0.030 (10) | 0.163 ± 0.026 (10) | 0.164 ± 0.020 (15) | 0.156 ± 0.014 (35) |

**Table 5b.**
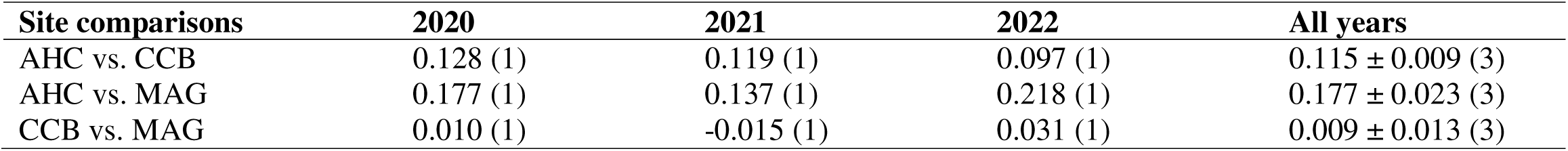
Pairwise *F_ST_* means and standard error for gametophytes in 2020, 2021, 2022, and across all years. Numbers in parentheses are the number of pairwise comparisons. Fixed sites (AHC, CCB, MAG) are characterized by thalli fixed to hard substratum via a holdfast

**Figure 7.**
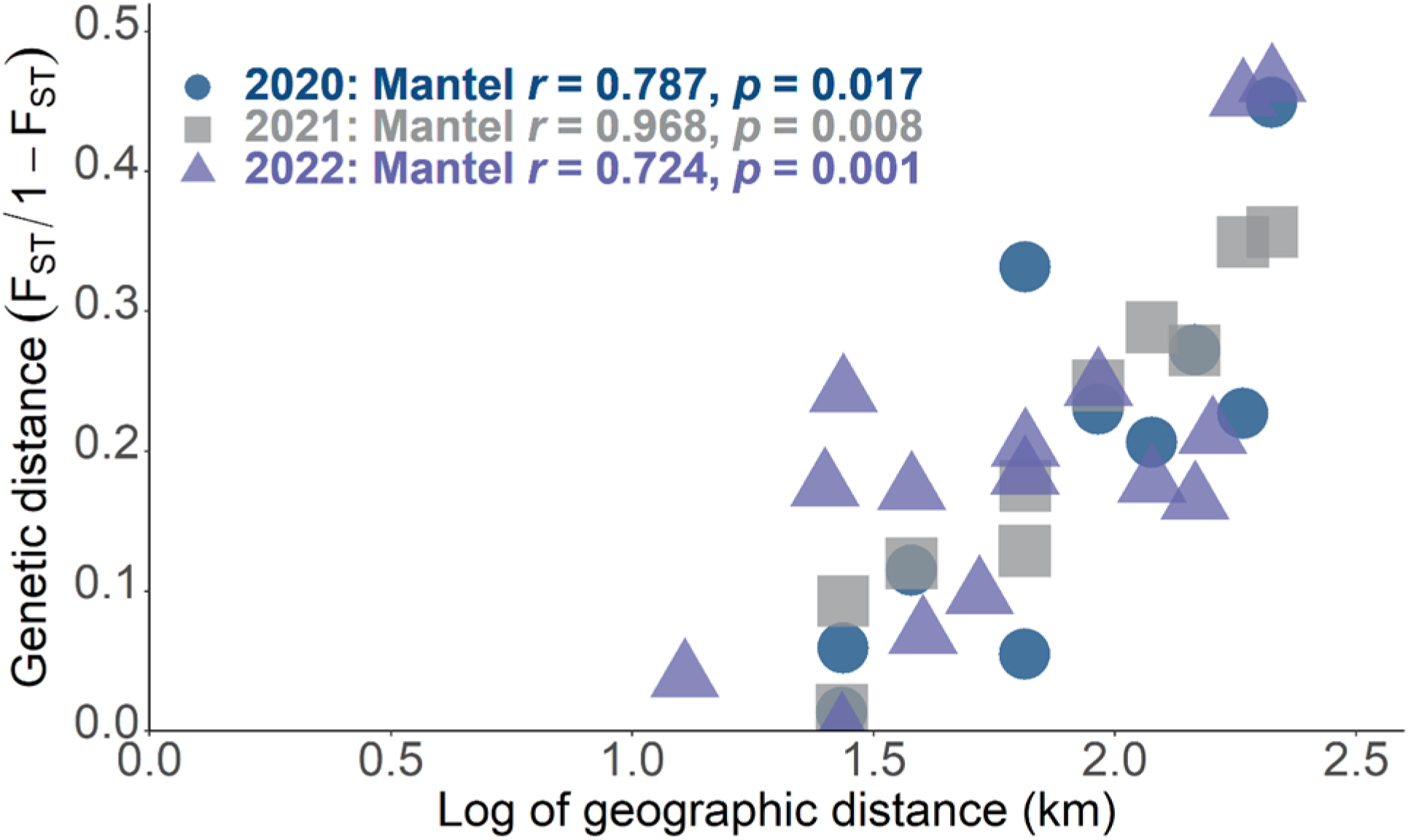
Genetic differentiation of *Gracilaria vermiculophylla* sites along the Delmarva Peninsula. Pairwise genetic distances, measured by allele identity (*F_ST_* / 1 - *F_ST_*) are plotted against pairwise geographic distances (km) for each coastline. For pairwise *F_ST_* for each year sampled, all slopes were significantly different from zero (2020: y = 0.329x - 0.425; 2021: y = 0.340x - 0.438; 2022: y = 0.247x - 0.246).

There were no differences in the pairwise genetic differentiation (*F_ST_*) when calculated between fixed gametophytes (Table S9i-p) or fixed tetrasporophytes within the same year or between years (Table 5b; within year: *W* = 30, *p* = 0.377; one year between sampling: *t_9.96_* = - 0.37, *p* = 0.729; two years between sampling: *t_3.73_* = -0.02, *p* = 0.985). There were also no differences when comparing the same site across years (Table 5b, Table S9i-p; *t_15.24_* = -2.83, *p* = 0.013).

*F_ST_* between fixed and free-living thalli were significantly higher than among fixed and among free-living thalli (H = 20.96, *p* < 0.001; Figure 8), indicating a hierarchical genetic differentiation between habitat type (e.g., hard versus soft bottom), then between sites, and finally between years. An analysis of the genetic differentiation dynamics (Figure 9) indicates that RED 2020 tended to spread into WAC 2021 and MAG 2021. RED 2021 still tended to spread into MAG 2022, but WAC 2021 spread into RED 2022. CCB both in 2020 and 2021 tended to spread into MAG 2021 and 2022. AHC 2020 tended to spread into CCB 2021, but CCB 2021 tended to spread into AHC 2022.

**Figure 8.**
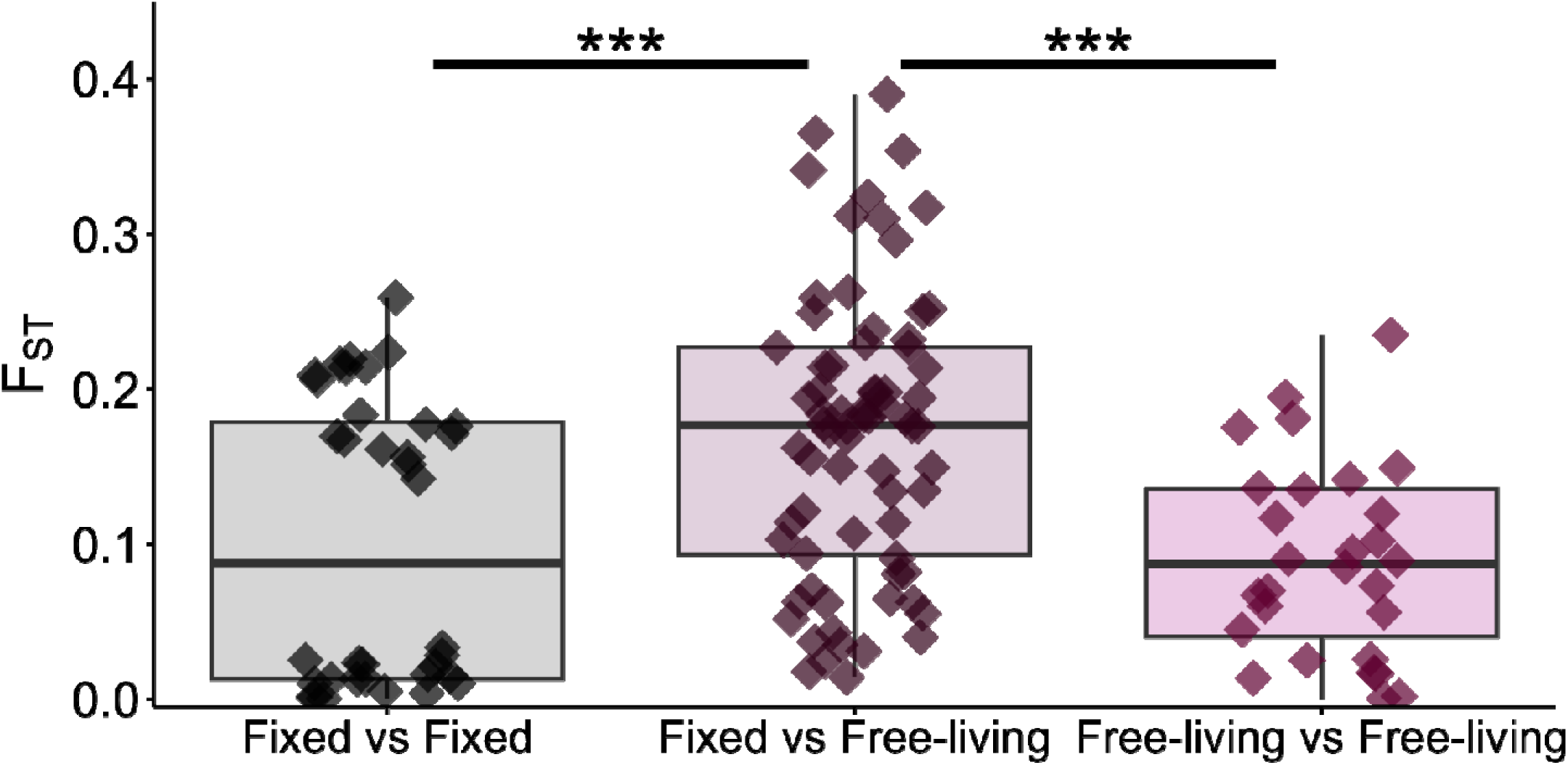
Distributions of *F_ST_* between pairs of hard bottom sites with fixed thalli, pairs of soft bottom sites with free-living thalli, and pairs of hard and soft bottom sites with fixed and free-living thalli, respectively. Kruskall-Wallis’ test indicated significant differences between at least two distributions. Based on Conover’s test of multiple comparisons, there were significantly different distributions between pairs of hard bottom sites with fixed thalli vs pairs of hard and soft bottom sites with fixed and free-living thalli (*p* < 0.001) and pairs of soft bottom sites with free-living thalli vs pairs of hard and soft bottom sites with fixed and free-living thalli (*p* < 0.001), pictured as three stars on the figure. There was no difference between fixed vs fixed and free-living vs free-living (*p* = 0.651).

**Figure 9.**
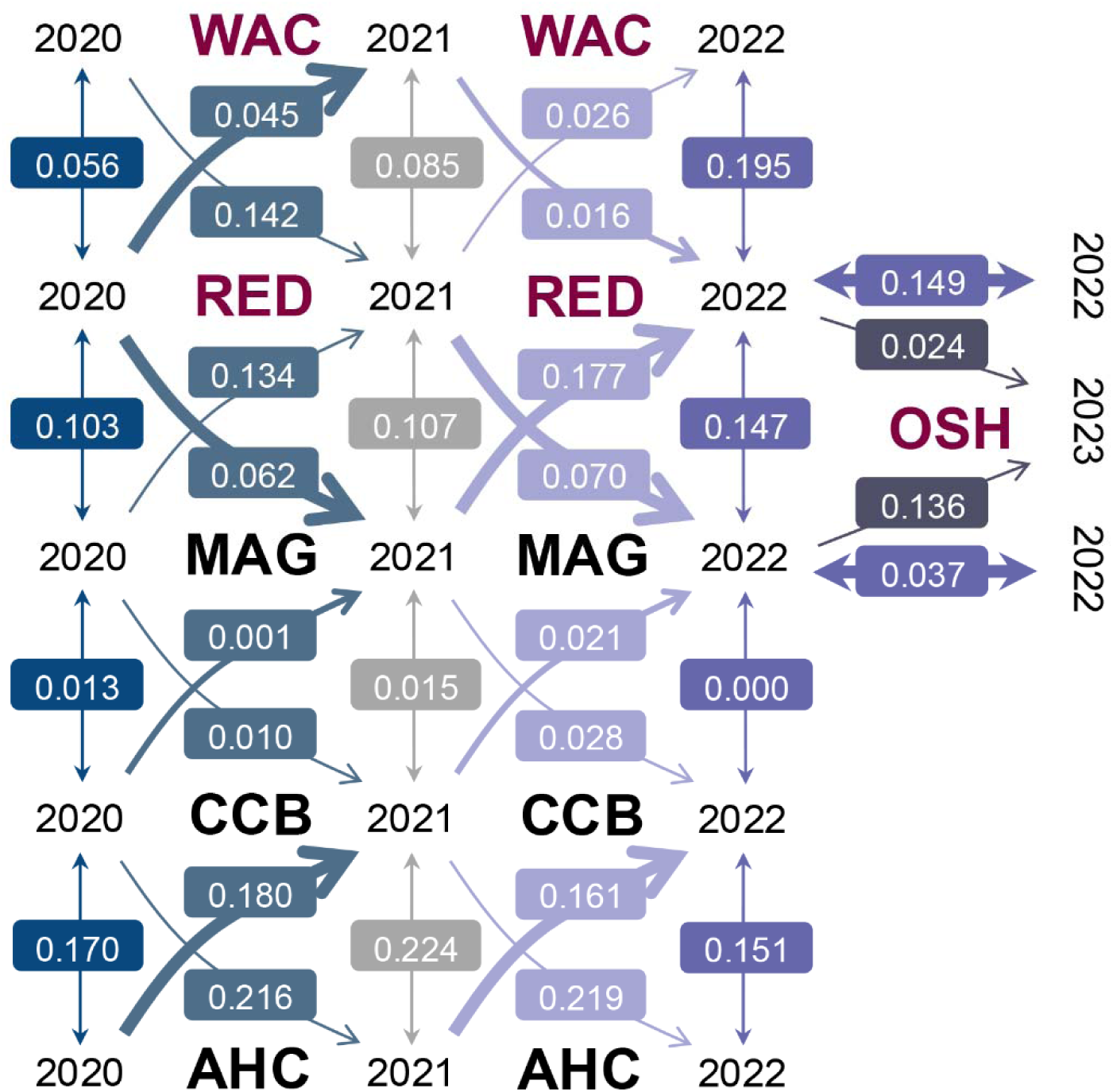
Dynamics of the genetic differentiation (*F_ST_*) between sites and years. The *F_ST_* and the thickness of the arrow between sites and years indicate direction of gene flow. By comparing the respective genetic differentiations between two sites from one year to another, we obtained the likely direction of gene flow. For example, *F_ST_* from RED 2020 to WAC 2021 was 0.045 (low) while *F_ST_* from WAC 2020 to RED 2021 was 0.142 (intermediate), showing that the genetic diversity in RED 2020 spread into WAC 2021 rather than the opposite direction. By comparing genetic differentiation between two sites a same year along the years indicated if genetic structure increased, was stable, or decreased between sites. For example, genetic differentiation increased from 2020 to 2022 between RED and WAC (0.056, 0.085, 0.195).

Sites grouped together generally on the minimum spanning tree of genetic distances between pairs of individuals, with MAG and RED at the center of the network (Figure 10). We found two types of populations: Within AHC, CCB and MAG, all thalli sampled locally belong to the same genetic background, but each year appeared as being an independent sample from a core population sharing the same locally coherent ancestralities (Figure 11a-c). In AHC, CCB and MAG, very distinct ancestralities structured by years may indicate fast renewal of populations in the place that were sampled, which is what is expected when sampling the perturbated edges of a larger population. By contrast, successive years at OSH and RED structured as being subsamples of the previous core year, radiating around the individuals of the previous year (Figure 11d,e). WAC pattern showed ancestralities structured by year with some individuals descending from previous years, in the terminal part of the network branches (Figure 11f).

**Figure 10.**
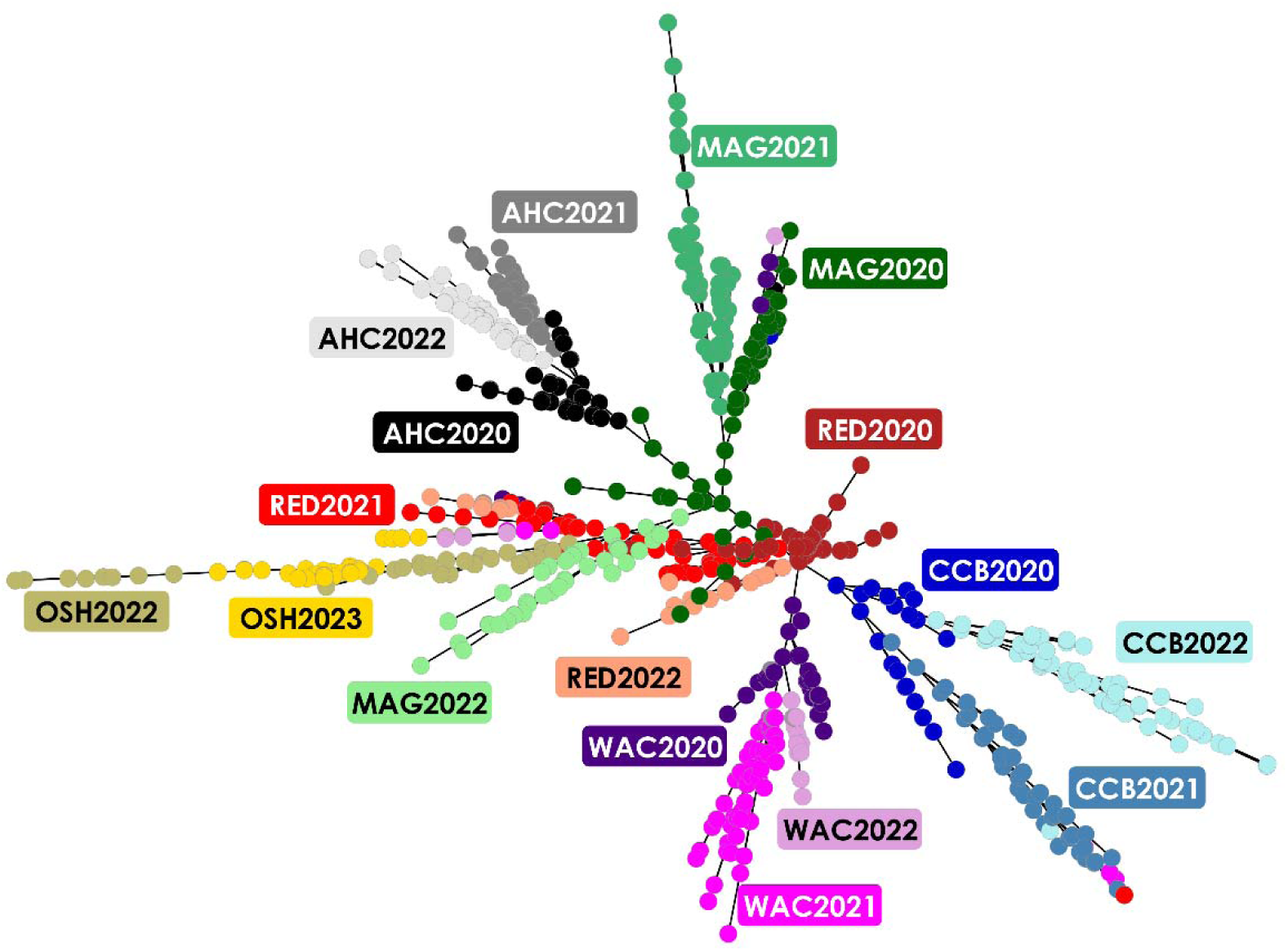
Minimum spanning tree illustrating the genetic relationships among thalli. Points are colored by site and year. AHC is shown in shades of gray, CCB in shades of blue, MAG in shades of green, OSH in shades of gold, RED in shades of red, and WAC in shades of violet. Distance following connection between two individuals (points) is proportional to their shared-in-state ancestrality. The network indicated individuals hierarchically structures as sites and years, with few migrants. Groups of individuals in rosary and rosette packs indicated clonal reproduction.

**Figure 11.**
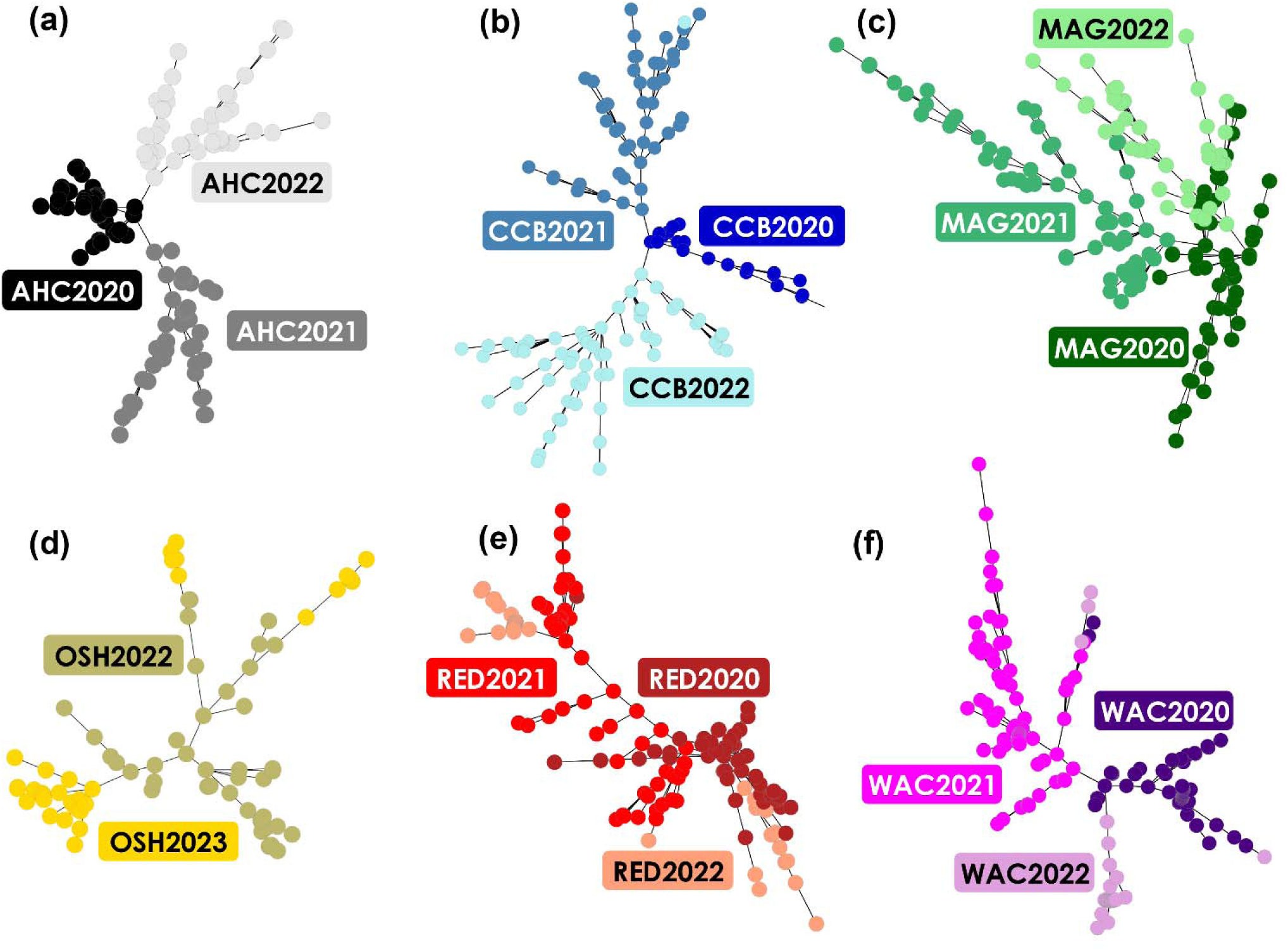
Minimum spanning tree illustrating the genetic relationships among thalli for (a) Ape Hole Creek (AHC) is shown in shades of gray, (b) Cape Charles Beach (CCB) in shades of blue, (c) Magotha Road (MAG) in shades of green, (d) Hillcrest (OSH) in shades of gold, (e) Fowling Point (RED) in shades of red, and (f) Upper Haul Over (WAC) in shades of violet. Points are colored by year.

## DISCUSSION

We hypothesized that fixed sites would be characterized by intergametophytic selfing due to bottlenecks associated with the invasion, while tetrasporophytic dominated free-living sites would be characterized by clonal reproduction with varying clonal rates. Overall, we found clear signs of clonality and selfing across sites. There was high power at all sites and time points, as overall *pidu* was always much less than 0.001 for tetrasporophytes and 0.006 for gametophytes. However, the probability of identity between sibs (*pids*) indicated that clonal processes can be difficult to disentangle from selfing processes, which is expected in populations combining autogamy and clonal reproduction (Stoeckel et al. 2024b). When we ranked sites according to clonality, soft bottom sites with free-living thalli were highly clonal with low genotypic diversity and dominated by tetrasporophytes. Hard bottom sites with fixed gametophytes and tetrasporophytes ranked from intermediate clonality to high clonality, depending on the demographic history of the site (i.e. frequent extirpation events versus stable populations).

However, due to the presence of male gametophytes, female gametophytes, and tetrasporophytes, the clonal rates at these sites should be interpreted with some caution and rather as a legacy of clonality across the genome further influenced by intergametophytic selfing. Genetic diversity and ancestralities were structured by habitat type (e.g., hard versus soft bottom), then by site, and finally by year, where there were patterns of isolation by distance.

Below, we discuss the eco-evolutionary consequences of reproductive mode variation, genetic structure, and ploidy bias observed in *G. vermiculophylla* along the Delmarva Peninsula.

### Reproductive mode variation at hard bottom sites with fixed thalli

In the sites we studied along the Delmarva Peninsula, there appears to be a complicated legacy of thallus fragmentation followed by likely migration into hard bottom sites. There is a mixture of intergametophytic selfing and outcrossing in differing levels across the genome and across years based on the demographic history of the site. *Gracilaria vermiculophylla* populations in the native range have high genetic diversity and positive *F_IS_* values consistent with intergametophytic selfing (Krueger-Hadfield et al., 2017). However, temporal genotyping is required to more accurately describe partial clonality by distinguishing between selfing and clonal reproduction as well as quantitatively infer their respective rates (Becheler et al., 2017).

We predicted intergametophytic selfing would prevail due to low gamete dispersal (Searles, 1980) and bottlenecks from the invasion (Flanagan et al., 2021; Krueger-Hadfield et al., 2017). We found no differences in *F_ST_* between phases regardless of site or year, suggesting regular gene flow between phases. The patterns we observed in *G. vermiculophylla* are comparable to the *F_ST_* between phases measured in other algal species for which allele frequencies were calculated in both the diploid and haploid subpopulations (see Krueger-Hadfield et al., 2021a).

Each of our three hard bottom sites had unique patterns of reproductive mode variation likely driven by overall ecology, stability of substrate, and environmental conditions. The thalli at MAG are found on the seaside of the Delmarva and grow on shell hash and wood pilings. This site has a more complex interaction with nearby mudflat, soft bottom habitats. CCB is the most stable hard bottom site, as thalli are fixed to jetties by holdfasts. This site is also the most like other rocky shore sites studied in *Gracilaria* spp. (e.g., Engel et al., 2004; Guillemin et al., 2013) and other red algae (see Krueger-Hadfield et al., 2021a). AHC, by contrast, is more like MAG in terms of the hard substrate to which thalli are fixed, including shell hash and wood pilings. Yet, thalli at this site are also found in consistently lower salinity as compared to all other sites included in our study. Moreover, previous data has suggested that thalli at this site are near their physiological limit for tolerating low salinity (see Krueger Hadfield & Ryan, 2020).

The fixed thalli in hard bottom habitats in *G. vermiculophylla* along the Delmarva appear to be undergoing varying rates of intergametophytic selfing. Potential technical causes of high *F_IS_* values, such as null alleles, do not contribute to the positive inflation of *F_IS_* in our study.

While we cannot discount spatial substructuring entirely as we did not sample following a hierarchical method, our observations of other genetic signatures in the hard bottom sites suggest fertilization regularly occurs between gametophytes sharing the same tetrasporophytic parent. At MAG and CCB, we observed a decline in outcrossing and increased selfing over the course of the sampling years. At AHC, we likely were sampling the edge of a large, panmictic population, but also sampled a small number of parents that sired the next generation, which confirms our hypothesis that fertilization can occur over small spatial scales. A combination of selfing and perhaps genotypic resolution based on our nine microsatellites made it difficult to disentangle clonal processes from selfing at AHC. Yet, it is unclear at present if intergametophytic selfing is more common in general in *G. vermiculophylla* as compared to observations in *G. gracilis*, where the populations studied were outcrossing (Engel et al., 2004). Krueger-Hadfield et al. (2016) observed positive *F_IS_* values in the sites sampled in the native range which could be interpreted as evidence of intergametophytic selfing. Yet, future efforts will need to sample populations in the native range to determine if selfing is common in *G. vermiculophylla* or if this is a consequence of the invasion in the non-native range. Moreover, paternity analyses would enable the direct characterization of the mating mode (i.e., selfing to outcrossing) as we can calculate the relatedness of male and female pairs, rather than relying on indirect measures such as *F_IS_* (Engel et al., 1999; Krueger-Hadfield et al., 2015). For example, in *Chondrus crispus*, male gametophytes were not only highly related to each other but also the female gametophyte that they fertilized (Krueger-Hadfield et al., 2015). Given that microspatial scale structure influence patterns of genetic structure in intertidal zones, paternity analyses are crucial to better resolve the mechanisms underlying the genetic patterns we have observed (see discussion in Krueger-Hadfield et al., 2013).

We did observe repeated genotypes at all three hard bottom sites. Free-living thalli are rare at these sites, and even if observed, we did not sample a free-living thallus. Moreover, *Gracilaria vermiculophylla* thalli do not reattach to hard substrate, possibility contributing to the re-sampling of ramets of the same genet following detachment and fragmentation as observed in other red algae (e.g., *Plocamium* spp., Heiser et al., 2023 or *Chondria tumulosa*, Williams et al., 2024). Re-encountered genotypes likely arise from a range of processes. Even though we did sometimes remove the entire holdfast, particularly at MAG or AHC, we sampled in the same area at each site at each sampling time. On more stable substrata, such as wooden pilings at MAG or AHC or the jetty at CCB, it is possible we re-sampled new upright thalli from the same holdfast. For repeated gametophytes, we have limited resolution because gametophytes are haploid and only have a single allele at each locus, thus are more likely to share an MLG despite being produced through independent meiotic and fertilization events. Previous studies have documented many more repeated gametophytic genotypes as compared to tetrasporophytes in *G. vermiculophylla* (Lees et al., 2018) and in *G. chilensis* (Guillemin et al., 2008). For repeated tetrasporophytes, it is also possible that repeated genotypes are the product of cystocarpic reproduction, whereby multiple tetrasporophytes share the same genotype because they were from the same fertilization event and released from the same carposporophyte. This was the expectation for Florideophyte red algae (see discussions in Engel et al., 2004; Krueger-Hadfield et al., 2011, 2013) as Searles (1980) hypothesis about the evolution of the carposporophyte phase centered on the rarity of fertilization events. Yet, there is very little evidence of tetrasporophytes sharing the same genotype in natural populations of *G. gracilis* (Engel et al., 2004) or *Chondrus crispus* (Krueger-Hadfield et al., 2013). Krueger-Hadfield et al. (2016) did not find many repeated tetrasporophytic genotypes in the native range for *G. vermiculophylla*. Thus, it is unclear if cystocarpic reproduction is perhaps greater in the non-native range and will require subsequent sampling. Increasing the number of loci will be critical to distinguish genotypes and resolve patterns of genotypic re-encounters, particularly across hard bottom sites with fixed thalli.

### Reproductive mode variation at soft bottom sites with free-living thalli

Consistent with previous surveys (Krueger-Hadfield et al., 2016, 2017), soft bottom sites were dominated by free-living tetrasporophytes and accompanied by signatures of clonal reproduction from thallus fragmentation. While we may have expected less compact ancestralities due to migration and mixing across soft bottom sites, the minimum spanning tree did not indicate this pattern. The combination of single time points and temporal genotyping revealed that high rates of clonality characterize soft bottom habitats. Moreover, much like our observations at the hard bottom sites, OSH, RED, and WAC each have unique environmental and ecological conditions that likely drive patterns of reproductive mode variation.

All three of the soft bottom sites are in the coastal bays of the Delmarva seaside and are connected by channels where free-living thalli could be dispersed. All three sites also exhibit fluctuations in biomass. The sites WAC and RED are the most similar in terms of size and distribution of *Gracilaria vermiculophylla*, most of which are anchored by *Diopatra cuprea*.

However, the two sites differed in important ways. We have monitored thalli at WAC consistently since 2014 (see Ross & Snyder (2024) and references therein) and find thalli year-round, though there are changes in biomass. This was also the site with the highest clonal ranking, and we consistently observed repeated genotypes both within and between years. By contrast, thalli at RED had variable abundance throughout the phenology survey, including one time point where thalli were entirely absent (Krueger-Hadfield et al., 2025b). It is likely the population size at RED is relatively small, with biomass declining over time and, notably, disappearing entirely from the mudflat during a survey in March 2025 (S.A. Krueger-Hadfield, *personal observation*). RED was characterized by an intermediate clonal rate. It is unknown where thalli originate here, especially after they are locally extirpated. However, there was evidence of thalli spreading from WAC to RED and vice versa. It is also likely that thalli are spreading from additional, unsampled sites. More detailed sampling in future studies will be essential to better track movement throughout the coastal bays.

In comparison to WAC and RED, the *Gracilaria vermiculophylla* thalli at OSH are found in a much narrower band, characterized by oyster reefs to the east and a channel to the west.

Thalli at this site were characterized by intermediate clonality, with a background signal of selfing. The origin of these tetrasporophytes is unclear, although nearby hard bottom sites, such as at MAG, may contribute new thalli that are the product of selfing. More detailed sampling is needed to clarify these dynamics and identify potential source sites.

### Genetic structure along the Delmarva

Genetic differentiation as measured by allele frequencies (*F_ST_*and ploidy-free *rho*) increased with geographic distance. Previous studies on *Gracilaria vermiculophylla* in this region have shown isolation by distance (Krueger-Hadfield et al., 2017). In the native range, there was also a high degree of genetic differentiation among sites that were in close proximity (∼10 to 50 km; Krueger-Hadfield et al., 2017). In *G. chilensis*, genetic differentiation was observed among populations from both wild and cultivated stands (Guillemin et al., 2008), an analog to our hard and soft bottom sites in *G. vermiculophylla*. Indeed, genetic differentiation was lowest between WAC, RED, and OSH, indicating potential thalli movement among these sites.

Overall, genetic diversity was structured first by habitat type, then by site, and finally by year. Curiously, thalli from RED tended to spread into the other soft bottom site WAC and the hard bottom site MAG. Thalli from WAC also tended to spread into RED, suggesting ongoing genetic connectivity between sites. Our genetic data also hint that thalli from CCB spread into MAG. However, it is not known how thalli move between fixed sites, such as CCB and MAG that are found on the bayside and seaside, respectively. Our genetic data suggest thalli from AHC have spread into CCB. The sites at AHC and CCB are located approximately 120 km apart and we currently do not have good distribution data for the bayside to determine if there are *G. vermiculophylla* populations found contiguously from CCB to AHC. While we have observed thalli at AHC on small pieces of shell hash moving with tidal rhythms (S.A. Krueger-Hadfield and W.H. Ryan, *personal observations*), this is unlikely to contribute to long distance dispersal.

### Substrate or invasion? Tetrasporophytic bias in Gracilaria vermiculophylla

Differences in mortality and fecundity rates may alter the ratio of gametophytes to sporophytes (Thornber, 2006). Similarly, fertilization rates and the number of carpospores produced have been shown to influence the phase ratio, especially at low fertilization rates (Fierst et al., 2005). Colonization events can also lead to phase bias, depending on which phase was initially introduced (Krueger-Hadfield, 2020). While sporophytic dominance has been observed in the Gracilariales (Kain & Destombe, 1995), the Ceramiales (Cecere et al., 2000), certain species of *Ulva* (Du et al., 2019), and brown algae, such as *Dictyota dichotoma* or *Ectocarpus crouaniorum* (Couceiro et al., 2015; Steen et al., 2019), other haploid-diploid algae can be gametophyte biased, such as observed in *Chondrus crispus* (see review in Collén et al., 2014) or *C. verrucosus* (Bellgrove & Aoki, 2008).

In our study, though gametophytes were common at hard bottom sites, sites were significantly tetrasporophyte biased. If thalli drifting into soft bottom sites continually arise from hard bottom sites, there would be more tetrasporophytes arriving based on observed phase ratios. However, this does not explain the complete absence of gametophytes in soft bottom habitats.

Previous studies on *Gracilaria vermiculophylla* have proposed several reasons why tetrasporophytes may dominate and this could apply to both habitat types. Gametophytes have been shown to have greater protein content and require less tensile force to detach a branch from the thallus, possibly translating into differences in herbivory based on tissue quality (Lees et al., 2018) and rate of fragmentation in free-floating conditions. In laboratory experiments, gametophytes suffer higher mortality rates across treatment conditions (Krueger-Hadfield & Ryan, 2020). In *G. chilensis*, tetrasporophytes may dominate because thalli are more robust to environmental stress and exhibit faster growth compared to gametophytes (Guillemin et al., 2013). Future studies should continue to investigate phenotypic differences between gametophytes and tetrasporophytes, including growth, survival, chemical composition, chemical defense, and biomechanical properties.

Previous work in *Gracilaria vermiculophylla* focused more on the comparison of native versus non-native populations (e.g., Bippus et al., 2018; Krueger-Hadfield et al., 2016; Murren et al., 2022; Sotka et al., 2018). Though we observed *G. vermiculophylla* thalli in soft bottom sites in the native range (see as an example, the site Futtu in Krueger-Hadfield et al., 2017), the preponderance of sites sampled were in hard bottom habitats, ranging from rocky shores to mudflat-type habitats with abundant hard substrate. By contrast, many sites in the non-native range were soft bottom without abundant hard substrate, with the largest numbers of sites sampled in the mid-Atlantic region south to the Carolinas along the US eastern seaboard (see Krueger-Hadfield et al., 2017, Sotka et al., 2018). As such, early work on the reproductive mode contrasted native versus non-native in the shift from sexual to clonal reproduction, but did not directly address the role of substrate type (Krueger-Hadfield et al., 2016). While the lability in the reproductive mode no doubt facilitated the invasion from Japan (e.g., Baker’s Law, Baker, 1955; Pannell et al., 2015; see also invasion history in Krueger-Hadfield et al., 2017), our data confirm that the shift toward clonal reproduction is largely driven to the substratum availability. Thus, the invasion of soft bottom habitats leads to the same consequences as observed in earlier native versus non-native comparisons (Krueger-Hadfield et al., 2016). Subsequent surveys, specifically along the westerns coast of the United States (Krueger-Hadfield et al., 2018), have revealed many more hard bottom habitats inhabited by *G. vermiculophylla*. Based on the complex interaction between soft and hard bottom populations in the current study, we propose that the clonal thallus fragmentation likely enhanced colonization success throughout the Northern Hemisphere and we expect to observe a complex legacy of clonality in hard bottom habitats throughout the non-native range as we have documented here along the Delmarva Peninsula. However, we expect that colonization of soft bottom habitats will result in shifts in both the prevailing reproductive mode as well as the phase ratio, regardless of site location (i.e., native or non-native). Future sampling is required to determine if colonization of soft bottom habitats is proportionally more common in the non-native range and to determine if similar population genetic dynamics occur in the native range.

### Conclusion and future directions

Our study was rooted in natural history observations that greatly enhanced our ability to interpret site-specific genetic patterns. Soft bottom sites showed tetrasporophytic dominance and high rates of clonality whereas hard bottom sites were more sexual. Though marginally non-significant there was a trend in which clonal rates increased over the sampling dates, while their selfing rates remained unchanged. This variation in population genetic indices warrants further investigation to identify potential underlying environmental factors, such as changes in temperature, that may explain such reproductive patterns. We also found repeated genotypes within and between years, which may be the result of sampling the same genet twice, an inability of the markers to distinguish among genotypes, or the product of cystocarp reproduction. Future studies should further investigate the spatial distribution of repeated genotypes within sites, particularly hard bottom sites.

As climate change and anthropogenic disturbances intensify, natural experiments develop if we track the distribution, abundance, and evolution of genetic traits in species’ responses. The *Gracilaria vermiculophylla* invasion provides an opportunity to investigate the evolution of biphasic life cycles and further understand reproductive mode diversity in natural populations.

Understanding the reproductive mode, especially in haploid-diploid taxa for which less information is readily available, is crucial for predicting how populations are likely to respond to environmental change.

## SUPPLEMENTAL TABLES

See supplemental Excel file

## SUPPLEMENTAL FIGURES

### ACKNOWLEDGEMENTS

We thank R. Snyder, S. Fate, P.G. Ross, J. Lewis, and E. Smith for field help and logistics at the Virginia Institute of Marine Science Eastern Shore Laboratory (VIMS ESL). We thank B. Thornton for help with July 2022 collections. We thank C. Amsler, J. McClintock, S. Watts, and R. Snyder for serving on for serving on the dissertation committee of A.P. Oetterer. This work was supported by an International Phycological Society Paul Silva Student Grant (to APO), start- up funds from the University of Alabama at Birmingham (UAB) College of Arts and Science (to SAKH), an NSF CAREER award (DEB-2141971 [UAB] and DEB-2436117 [VIMS|W&M] to SAKH), NSF OIA-1946412 (to SAKH), ANR-18-CE32-0001 (to SS and SAKH), and start-up funds from the Virginia Institute of Marine Science William & Mary (to SAKH). The Department of Biology at UAB provided logistical support. VIMS and VIMS ESL provided additional facilities and logistical support. A.P. Oetterer was supported by the UAB Blazer Fellowship. WHR was supported by the NIH IRACDA MERIT postdoctoral fellowship (NIH K12GM088010). S.A. Krueger-Hadfield was supported by the NSF (CAREER Award DEB- 2141971 and EAGER DEB-2113745).

## AUTHOR CONTRIBUTIONS

**Alexis P. Oetterer** Data curation (equal), Formal Analysis (equal), Funding acquisition (supporting), Investigation (equal), Visualization (equal), Writing – original draft (equal), Writing – review and editing (equal); **Solenn Stoeckel** Formal analysis (equal), Funding acquisition (supporting), Investigation (supporting), Software (lead), Visualization (supporting), Writing – original draft (supporting), Writing – review and editing (equal); **Will H. Ryan** Conceptualization (supporting), Investigation (supporting), Writing – review and editing (equal); **Stacy A. Krueger-Hadfield** Conceptualization (lead), Data curation (equal), Formal Analysis (equal), Funding acquisition (lead), Investigation (equal), Methodology (lead), Project Administration (lead), Resources (lead), Software (supporting), Supervision (lead), Visualization (equal), Writing – original draft (equal), Writing – review and editing (equal)

^1^ Angiosperms have an alternation of generations in which the sporophyte is dominant. The gametophyte is always retained on the sporophyte and is composed of only a few cells. Though the gametophytic phase is selectively important (Delph, 2019), we can consider angiosperms as ecologically diploid (see also Krueger-Hadfield et al., 2024).

---

**Figure S1.**
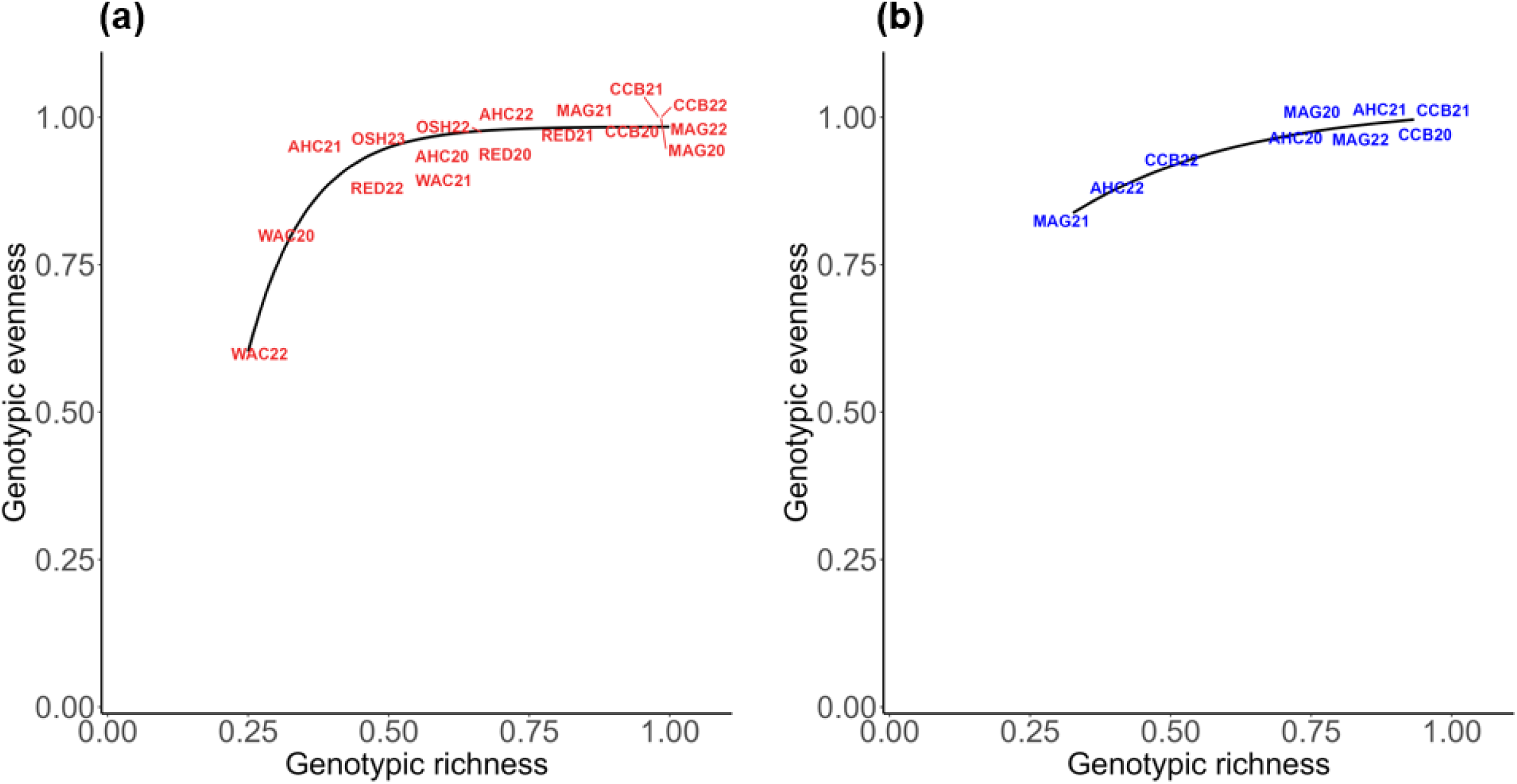
Reproductive mode variation shown as genotypic evenness (*D\**) versus genotypic richness (*R*). We fit an asymptotic regression for the (a) tetrasporophytes across all sites and years and (b) for the gametophytes across all hard bottom sites and years. Points are shown as site abbreviations and a two-digit year (20, 21, 22, 23).

**Figure S2.**
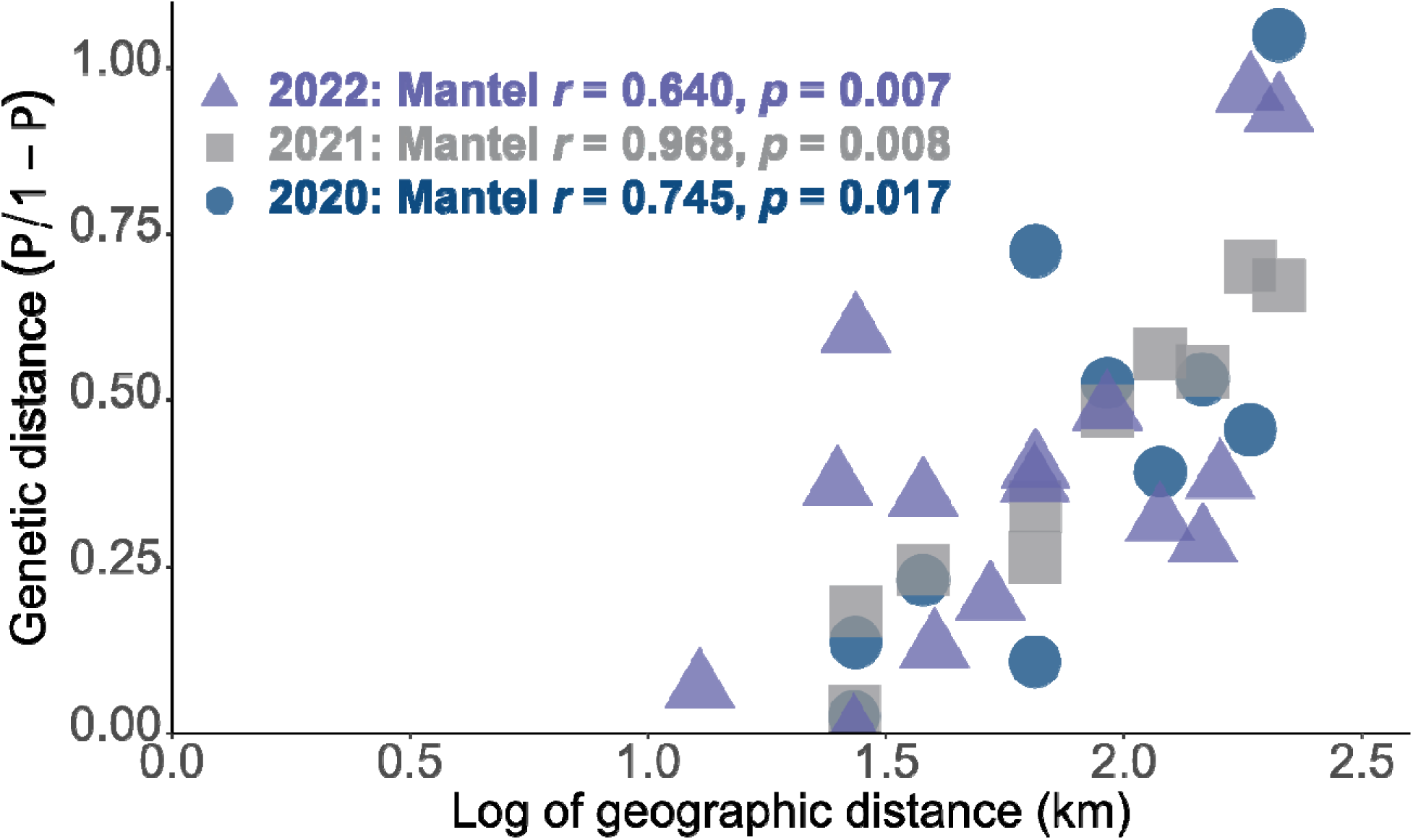
Genetic differentiation of *Gracilaria vermiculophylla* sites along the Delmarva Peninsula. Pairwise genetic distances, measured by allele size (*rho /* 1 *- rho*) are plotted against pairwise geographic distances (km) for each coastline. 2020: y = 0.712x - 0.926; 2021: y = 0.659x - 0.840; 2022: y = 0.464x - 0.432.

